# Antibiotic exposure dynamically generates a substantial number of heterogeneous persisters along a spectrum of tolerance

**DOI:** 10.64898/2026.01.30.702957

**Authors:** Yijie Deng, Douglas R. Beahm, Hannah E. Maurais, Kai Etheridge, Daniel Schultz, Rahul Sarpeshkar

**Affiliations:** Thayer School of Engineering, Dartmouth College, Hanover, NH 03755, USA; Department of Microbiology and Immunology, Geisel School of Medicine, Dartmouth College, Hanover, NH 03755, USA; Departments of Engineering, Physics, Microbiology and Immunology, Molecular and Systems Biology, Dartmouth College, Hanover, NH 03755, USA

**Keywords:** Antibiotic persistence, phenotypical heterogeneity, drug-induced persisters, persistence spectrum, kinetic modeling

## Abstract

Antibiotics are known to induce new persister cells during treatment, yet the inability to distinguish and quantify pre-existing versus drug-induced persisters has long obscured how antibiotics and genes shape persistence. Here, we develop a quantitative framework integrating kinetic modeling with serial-dilution time-kill (SDTK) assays to resolve persister population dynamics and accurately quantify both persister types. We show that antibiotic exposure dynamically generates a substantial number of persisters that are heterogeneous and distributed along a persistence spectrum. Across antibiotics, we uncover pronounced differences in rates of persister induction and elimination, with ampicillin inducing persisters at the highest rate and kanamycin at the lowest. Depending on dilution history, drug-induced persisters can dominate the persister pool. Our framework enables identification of genetic determinants specific to pre-existing and/or drug-induced persistence and reveals drug-dependent pre-existing persister fractions. Systematic sequential-drug treatments demonstrate that kanamycin persisters form the most tolerant subset, embedded within ciprofloxacin persisters that in turn are nested within the broader ampicillin persister subpopulation. Together, we propose a Drug-Induced Persistence-Spectrum (DIPS) model in which antibiotics differentially induce and select for persister subsets along a tolerance continuum.

## Main

The global crisis of antimicrobial resistance (AMR) is a major driver of antibiotic treatment failure ^1^, but non-heritable mechanisms such as antibiotic persistence also substantially undermine therapeutic efficacy, promote recurrent and chronic infections, and facilitate the evolution of antibiotic resistance ^2–5^. Antibiotic persistence arises from rare phenotypically tolerant cells, called “persisters”, which survive lethal antibiotic exposure without acquiring genetic resistance. These cells typically constitute <0.1% of bacterial populations, adopt metabolically slow or dormant states, and resuscitate after drug removal ^6,7^.

Although a myriad of different molecular pathways and stochastic mechanisms have been identified and reviewed for persister formation ^2,7–17^, their relative contributions, degree of independence, and hierarchical interactions remain widely debated. Persisters are commonly categorized as spontaneous persisters, which arise stochastically during growth and are extremely rare, and triggered persisters, which emerge in response to environmental stress and dominate the persister population ^10,16^.

The prevailing view of antibiotic persistence can be considered as the “pre-existing persister model,” which posits that nearly all persister cells exist before antibiotic exposure and that treatment merely reveals persisters without altering their numbers in time-kill assays ^18^. However, accumulating evidence suggests that antibiotics themselves can promote entry into persistence ^18–21^. Therefore, we propose a more general framework—the “drug-induced persister model”—in which antibiotic treatment not only reveals but also induces persisters. This new model accounts for both persisters that exist before antibiotic exposure (pre-existing persisters) and those generated during the treatment (drug-induced persisters).

Despite decades of research, the field still lacks a reliable methodology to distinguish the two persister subsets. This limitation has long prevented accurate quantification of treatment-emergent persisters and obscures how different antibiotics or genetic factors influence persister formation. Single-cell approaches yielded valuable insights into phenotypic heterogeneity and antibiotic persistence ^17,22,23^. However, these techniques remain constrained by the lack of definitive biomarkers for persister cells and insufficient resolution and throughput to reliably track these extremely rare persisters. This challenge extends beyond bacterial systems to drug-persistent yeasts and drug-tolerant persister cancer cells, where surviving populations may include both pre-existing persisters and cells that actively transition into tolerance during treatment ^24–27^. Across biological systems, the inability to discriminate these populations has hindered efforts to characterize the dynamic formation of persisters during treatment and to determine the roles of treatment-induced molecular reprogramming in persistence development. Further, this methodological gap obstructs the development of therapeutic strategies aimed at reducing treatment-induced persistence.

Here, we address this long-standing challenge by developing a quantitative kinetic framework that integrates mathematical modeling with a newly designed serial-dilution time–kill (SDTK) assay. This quantitative approach validates the more general “drug-induced persister model” and enables direct quantification of both pre-existing and drug-induced persisters within the same culture. While drug-dependent persister fractions show substantial heterogeneity in persistence, the relationship among persister subsets across antibiotics has remained unclear. By combing this framework with systematic sequential-drug-treatment experiments, we demonstrate that persistence is a dynamic, drug-induced process that generates heterogeneous persister subsets nested and distributed along a spectrum of tolerance. Our findings support a drug-induced-persistence-spectrum (DIPS) model that could reshape the conceptual foundation of antibiotic persistence and provide a generalizable strategy for dissecting treatment-induced phenotypic tolerance.

## Results

### Experimental design and quantification of the drug-induced persisters

We establish a simple yet powerful methodology to distinguish between the pre-existing-persister and drug-induced-persister models. We integrate biphasic kinetic modeling with an experimental strategy: the serial-dilution time-kill (SDTK) assay (Fig. 1a). In this assay, a single overnight seed culture is serially diluted (e.g., 100×, 1,000×, 10,000×) into fresh pre-warmed medium, and all resulting cultures are grown to the same optical density in exponential phase before antibiotic addition, followed by standard time-kill measurements. The pre-existing persisters in these cultures should scale proportionally to the serial dilution factor (e.g., 10x). Fitting experimental data to mathematical models allows us to distinguish between two models, determine the key parameters, and quantify both pre-existing and drug-induced persisters (Fig. 1b).

**Fig. 1.**
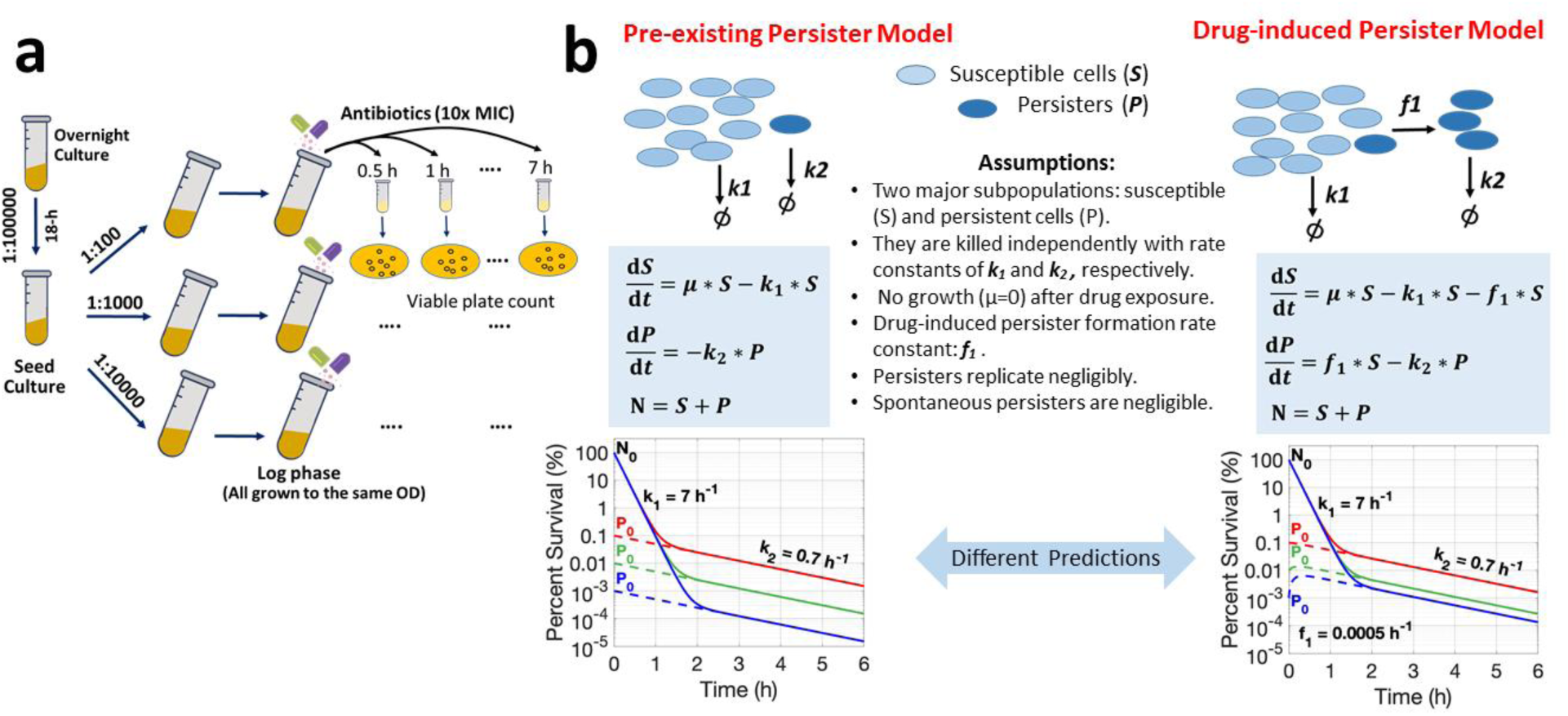
Methodology to distinguish two working models for antibiotic persistence. (a) Serial-dilution time-kill (SDTK) assay. Seed cultures are serially diluted in a fresh medium (e.g., 100x, 1000x, 10000x) and grown to an equal optical density at the exponential phase before exposure to antibiotics for standard time-kill assays. After drug removal, surviving cells are enumerated by counting colony-forming units (CFUs) using the viable plate count method. (b) Predictions of the pre-existing and drug-induced persister models. In the pre-existing model, all persisters exist prior to antibiotic exposure and are diluted proportionally with each serial dilution, resulting in evenly spaced secondary killing phases (10× dilution shown). In the drug-induced model, antibiotic stress induces additional persisters during treatment, causing the more diluted cultures—those with fewer pre-existing persisters—to show relatively elevated survival curves. Dashed curves denote the persister fraction, while solid curves denote the total population. Simulations were performed using biologically relevant parameters.

The rationale for this approach is based on two well-established observations. First, spontaneous persisters arise at extremely low frequencies during exponential growth ^10,16^, making their contribution negligible relative to the persisters originating from the stationary-phase inoculum. Second, persister cells, while metabolically active at low levels, replicate minimally upon re-inoculation into fresh medium during early exponential growth ^7,15,28,29^. Under these assumptions, the pre-existing persister model predicts that persister abundance should scale inversely with dilution: more diluted cultures contain proportionally fewer persisters, yielding evenly spaced secondary killing phases across serially diluted time–kill curves (Fig. 1b).

In contrast, the drug-induced persister model predicts a distinct and testable outcome: if antibiotics induce new persisters during treatment, then the more diluted cultures should show greater increases in relative survival because newly formed persisters constitute a larger proportion of the persister pool when the initial persister number is smaller. Thus, the divergent predictions (Fig. 1b) are uniquely resolved by the SDTK assays (see Methods for details).

We performed SDTK assays on *E. coli* treated with three classes of bactericidal antibiotics with distinct mechanisms of action: ampicillin (cell-wall synthesis inhibitor), ciprofloxacin (DNA replication inhibitor), and kanamycin (translation inhibitor). Across all three antibiotics, we observed a consistent pattern: instead of showing proportionally lower survival in serially diluted cultures (predicted by the pre-existing persister model), survival fractions were elevated with higher dilutions (Fig. 2a). Our results unambiguously demonstrates that antibiotic exposure, even at high doses (10× MIC), rapidly triggers the formation of persisters, consistent with the drug-induced persister model.

**Fig. 2.**
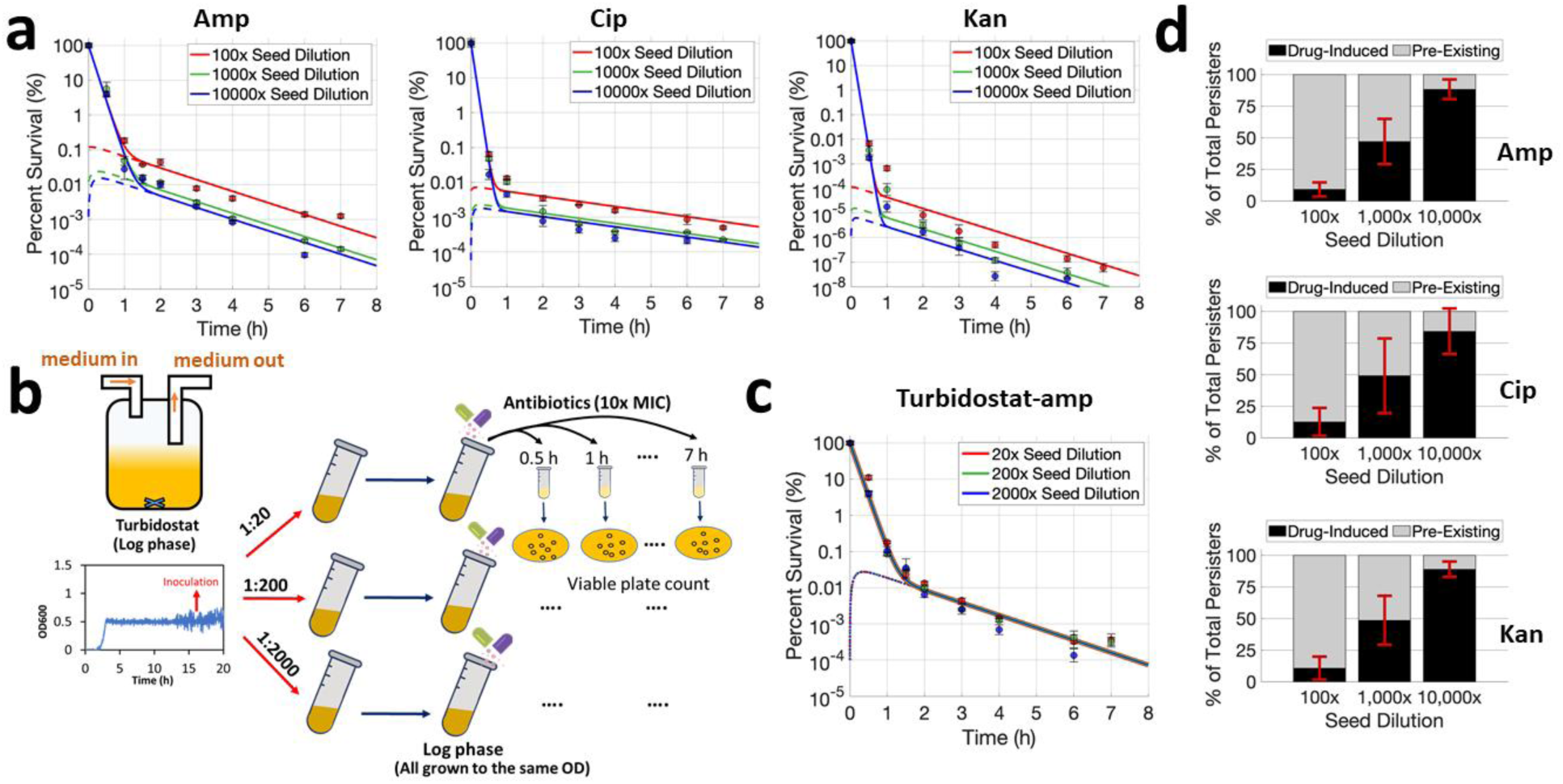
Experimental validation and quantification of drug-induced persister formation using SDTK assays. (A) SDTK assays of *E. coli* MG1655 treated with ampicillin (Amp, 80 µg/ml), ciprofloxacin (Cip, 1.25 µg/ml), or kanamycin (Kan, 320 µg/ml), each at 10× the minimal inhibitory concentration (MIC). Data were fitted using the drug-induced persister model. Shown are representative graphs from four replicates across two independent experiments. Dashed curves represent the persister subpopulation, whereas solid curves represent the total population. (b) Experimental setup for turbidostat-controlled growth followed by SDTK assays. Bacteria were maintained at constant optical density (OD₆₀₀ = 0.6) during exponential growth for approximately 12 h in the turbidostat, effectively minimizing persisters carried over from the stationary-phase inoculum. After serial dilution into pre-warmed fresh medium, the resulting cultures were re-grown to the same OD₆₀₀ values prior to ampicillin exposure. (c) SDTK assays of cultures derived from the turbidostatic experiments, with data fitted by the drug-induced persister model. (d) Relative fractions of drug-induced and pre-existing persisters within the total persister subpopulation for overnight batch experiments. Fractions were calculated from data collected 3 h after antibiotic exposure, at which point the proportions had reached steady states. Data are presented as means ± standard deviation (SD) from at least three independent experiments.

Turbidostatic experiments further validated the drug-induced persister model (Fig. 2b,c). In continuous exponential growth, persisters carried over from the stationary phase are diluted to negligible levels, such that persister cells must predominately arise from drug-induced formation regardless of initial inoculation density. Consistent with this prediction, all dilution series yielded nearly overlapping time–kill curves (Fig. 2c). These results confirm that antibiotic exposure alone can generate persisters.

An alternative explanation is that more-diluted cultures undergo longer exponential growth before reaching the same optical density, potentially generating more spontaneous persisters. However, this effect is partially offset by the proportionally smaller initial population size. To conclusively rule out this possibility, we developed a quantitative model incorporating spontaneous persister formation during exponential growth based on a previous framework ^16^ and simulated population dynamics from inoculation through 8 h after antibiotic exposure. Spontaneous persister formation had negligible effects on survival and failed to reproduce the SDTK patterns (Fig. S1), even when the spontaneous formation rate (*a₂*) was increased 100-fold above previously measured values ^16^ (Fig. S1f,g). A combined spontaneous and drug-induced model was similarly unnecessary (Fig. S1e). Similar results were observed in turbidostatic simulations (Fig. S2), which requires biologically implausible (200–650-fold) increases in *a₂* to approximate the data (Fig. S2g; Table S1). In contrast, simulations show that >99.8% of persisters arose during antibiotic exposure (Table S1), demonstrating that drug-induced persister formation alone explains the observed dynamics.

Combining kinetic modeling with SDTK experiments, we accurately quantified both pre-existing and drug-induced persisters within the same population—an otherwise extremely difficult task. Across all three antibiotics, a substantial fraction of persisters arose during antibiotic exposure. The relative contributions of the two persister types depended strongly on the dilution factor (i.e., initial number of pre-existing persisters). Drug-induced persisters comprised ∼10% of total persisters in 100-fold diluted cultures, ∼45% in 1,000-fold dilutions, and >80% in 10,000-fold dilutions (Fig. 2d). Although drug-induced persisters are generated in similar absolute numbers across dilutions, their relative contribution increases at higher dilutions because the initial pre-existing persister pool size is smaller. These results demonstrate that drug-induced persisters can dominate the persister subpopulation at high seed dilutions, challenging the prevailing view that pre-existing persisters overwhelmingly dictate the persister pool.

The dynamic nature of antibiotic persistence cautions against conventional persister measurements. First, persister frequencies are time-dependent such that sampling time strongly influences apparent persister frequencies. Second, because the persister pool comprises both pre-existing and drug-induced persisters, reporting a single persister fraction without distinguishing these subsets can confound comparisons and obscure gene or stress-response functions. Third, apparent persister levels depend on dilution history, with less-diluted cultures yielding higher values, necessitating careful control across conditions. Together, these considerations underscore the need to account for persister subtype, dilution history, and timing when quantifying and comparing persistence.

### The dynamics of antibiotic killing and persister formation depend on drugs

Our framework enables quantitative, side-by-side comparisons of antibiotic efficacy against both susceptible and persister subpopulations across antibiotics (all at 10× MICs). In addition, it provides a simple and robust means to measure persister-formation rates. Using this approach, we found that the rate at which persisters emerge during treatment is strongly drug-dependent, following the order ampicillin > ciprofloxacin > kanamycin (Fig 3a). This ranking indicates that ampicillin exposure induces persister formation more rapidly than ciprofloxacin or kanamycin exposure.

**Fig. 3.**
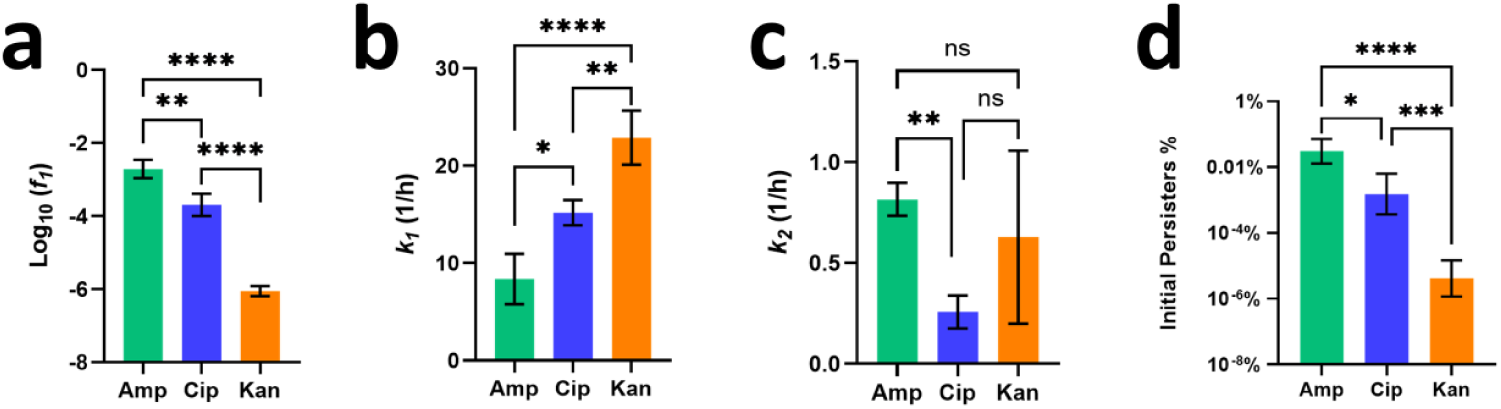
Persister formation and killing kinetics depend on antibiotic types. (a) Persister formation rates vary across antibiotics. Amp: ampicillin; Cip: ciprofloxacin; Kan: kanamycin. (b) Susceptible cells are killed with different rate constants (*k₁*) across three antibiotics. (c) Persisters are killed with distinct rate constants (*k₂*) across different antibiotics. (d) Pre-existing persister fractions (P₀%) in exponential-phase cultures before exposure to different drugs. Initial persister fractions (P₀%) were calculated by model fitting and are shown for the exponential-phase cultures derived from 1000-fold seed dilutions. Data are presented as means ± SD from at least three independent replicates. **** p < 0.0001; *** p < 0.001; ** p < 0.01; * P < 0.05; ns, not significant.

Drug efficacy varied substantially across antibiotics in terms of their killing activity against susceptible and persister subpopulations. Although ampicillin eliminates susceptible cells more slowly than ciprofloxacin and kanamycin (Fig. 3b), it kills persister cells faster than ciprofloxacin but not kanamycin (Fig. 3c). These results illustrate that the relative potency of an antibiotic against actively growing cells does not necessarily predict its effectiveness against persisters, emphasizing that persistence represents a distinct physiological state with drug-specific vulnerabilities.

We identified significant drug-dependent differences in pre-existing persister fractions. In exponential-phase cultures derived from a 1,000-fold diluted inoculum, the pre-existing persister fraction for ampicillin (∼0.03%) was more than an order of magnitude higher than that for ciprofloxacin, which in turn was more than two orders of magnitude higher than that for kanamycin (Fig. 3d). These results indicate that persister fractions depend strongly on the antibiotic used and are therefore comparable only within assays involving the same drug; cross-antibiotic comparisons of persister abundance should be interpreted with caution.

Our findings further reveal that persisters are not the same even before antibiotic exposure and possess drug-specific survival thresholds. A cell persistent to ampicillin is not necessarily persistent to ciprofloxacin or kanamycin, indicating that persister phenotypes are not homogeneous but instead reflect various metabolic or physiological states tailored to each antibiotic class ^30^. The substantially higher abundance of ampicillin persisters than ciprofloxacin or kanamycin persisters from the same culture (Fig 3d, Fig S4 b) indicates the possibility that ampicillin persisters are less persistent than the latter two and thus easier to form during drug exposure with higher *f_1_* (Fig. 3a) (more discussion below).

### Dissecting gene functions in the formation of pre-existing and drug-induced persisters

Quantitative separation of pre-existing persisters from those formed during drug exposure enables the mechanistic dissection of gene functions underlying persistence—specifically, whether a gene influences the formation of pre-existing persisters, drug-induced persisters, or both. Using the SDTK approach, we measured killing kinetics, persister fractions, and persister formation rates in wild-type and mutant strains lacking key regulators of the stringent response and stress adaptation. We specifically examined whether ppGpp (synthesized by RelA and SpoT) promotes the formation of pre-existing persisters during stationary phase and/or new persisters during antibiotic treatment, a distinction that is obscured in conventional time-kill assays. Additionally, inorganic polyphosphate (polyP) is a key regulator of bacterial metabolism and stress responses ^31,32^, yet its contribution to antibiotic persistence remains elusive. Therefore, we also evaluated the role of polyP, synthesized by polyphosphate kinase (PPK) in each of these processes. Representative SDTK assay time-kill curves for *Δppk* and *ΔrelAΔspoT* are shown in Fig. 4a,b.

**Fig. 4.**
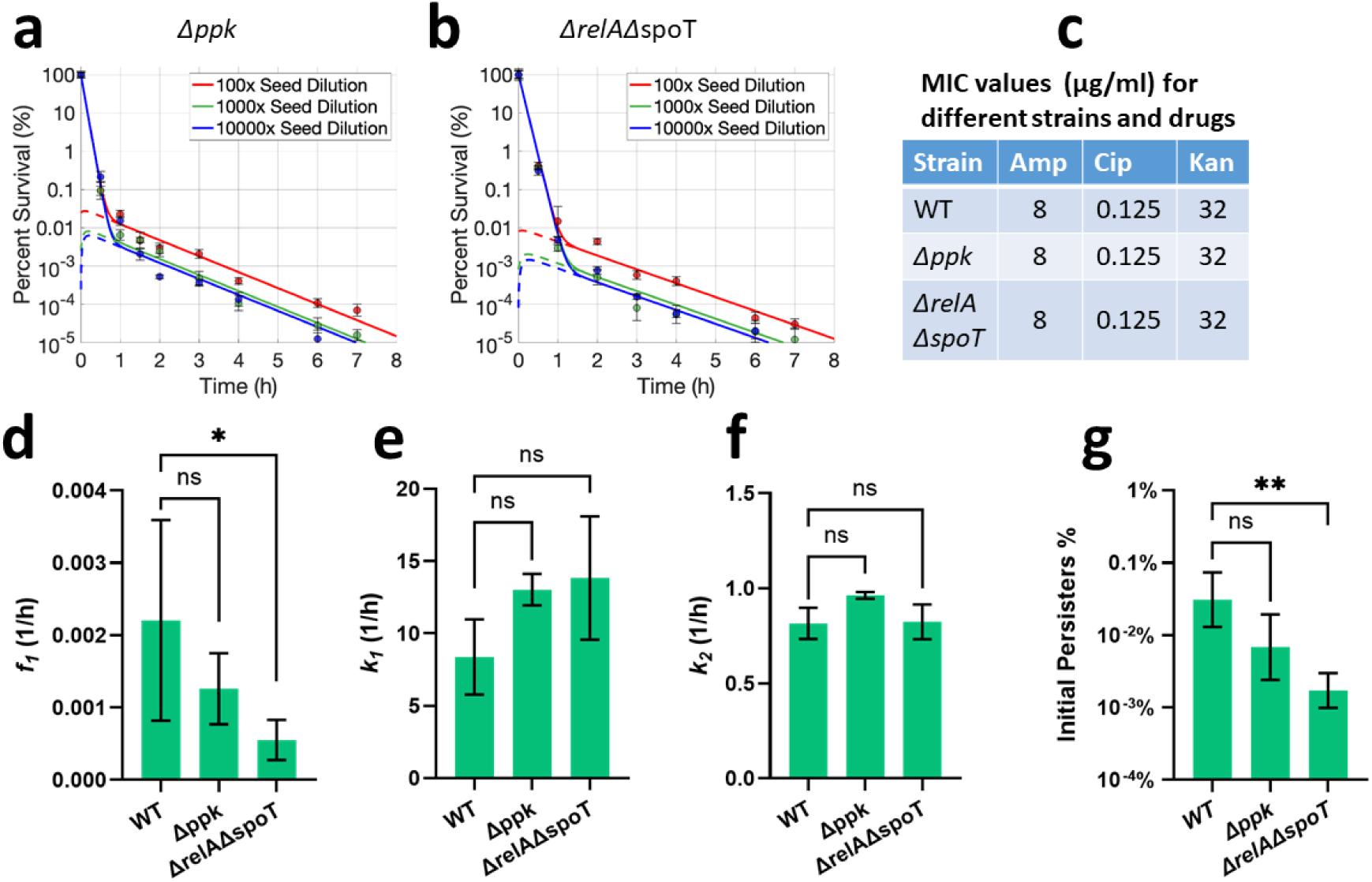
Roles of ppGpp and PPK in persister formation and antibiotic killing. (a, b) SDTK assays for *Δppk* and *ΔrelAΔspoT* strains exposed to ampicillin. *Δppk: gene knockout of polyphosphate kinase gene (ppk); ΔrelAΔspoT: ppGpp null strain.* Representative graphs from four replicates across two independent experiments are shown. Dashed curves represent the persister subpopulation, whereas solid curves represent the total population. (c) No difference in MIC values for wild-type and mutant strains among antibiotics tested. Amp: ampicillin; Cip: ciprofloxacin; Kan: kanamycin. (d-f) Comparing persister formation rates and killing rates across the wild-type and mutant strains treated with ampicillin. (g) Initial persister fractions (*P₀%*) of mutant strains treated with ampicillin across different strains. The fractions were calculated from the exponential-phase cultures seeded from 1000-fold dilutions. Data are presented with means ± SD with at least three independent experiments. ** p < 0.01; * p < 0.05; ns, not significant.

Although MIC values were unchanged in the *ΔrelAΔspoT* and *Δppk* strains compared to the wild-type strain (Fig. 4c), the *ΔrelAΔspoT* mutant exhibited substantially reduced persister-formation rates during ampicillin exposure (Fig. 4d), demonstrating that ppGpp is responsible for the generation of drug-induced persisters. Meanwhile, this mutant produced far fewer pre-existing persisters (Fig. 4g), indicating that ppGpp also increases persister formation during the stationary phase in addition to its role in forming persisters during drug exposure. The *ΔrelAΔspoT* strain’s strong defect in persister formation is consistent with the central role of ppGpp in metabolic downshifting and antibiotic persistence ^12^. The *Δppk* mutant consistently showed reduced persister formation rates and lower levels of pre-existing persisters compared to the wild-type strain (Fig. 4d,g), supporting a positive role for polyP in promoting antibiotic persistence, although these differences did not reach statistical significance. In addition to the altered persister dynamics, susceptible cells of both mutants appeared to be killed more rapidly (with larger *k_1_*), although the differences were not statistically significant (p > 0.05). In sum, while polyP plays a minor role in persistence, ppGpp strongly promotes persister formation during both the stationary phase and antibiotic treatment, presumably by slowing metabolism, arresting growth, and activating stress-response pathways such as the SOS and oxidative-stress responses^12,33,34^, collectively protecting cells from stress-induced damage.

### Dynamics of persister formation and antibiotic killing

A key advantage of our approach is that it resolves the population-level dynamics of persister formation without using specialized single-cell platforms. Analyzing 1000-fold diluted cultures as an example, our model shows that new persisters emerge rapidly during the earliest phase of antibiotic exposure, typically within the first 20 minutes, regardless of antibiotics (Fig. 5a). The rapid rise in persister numbers is more pronounced in cultures derived from 10,000-fold dilutions (Fig. S3a,b). This observation can be readily explained: pre-existing persisters are present at much lower levels in these cultures, so even a modest persister-formation rate (*f₁*) can generate a comparatively substantial number of new persisters from the large susceptible cell population within a small time window; as a result, newly-formed persisters quickly outnumber pre-existing persisters and dominate the persister subpopulation within 20 min (Fig. S3c). This timing is also consistent with reports that fluoroquinolones trigger immediate changes in the gene expression profile including the SOS response and DNA repair programs within 10-25 minutes ^19,35^, which can drive rapid induction of persisters ^19^.

**Fig 5.**
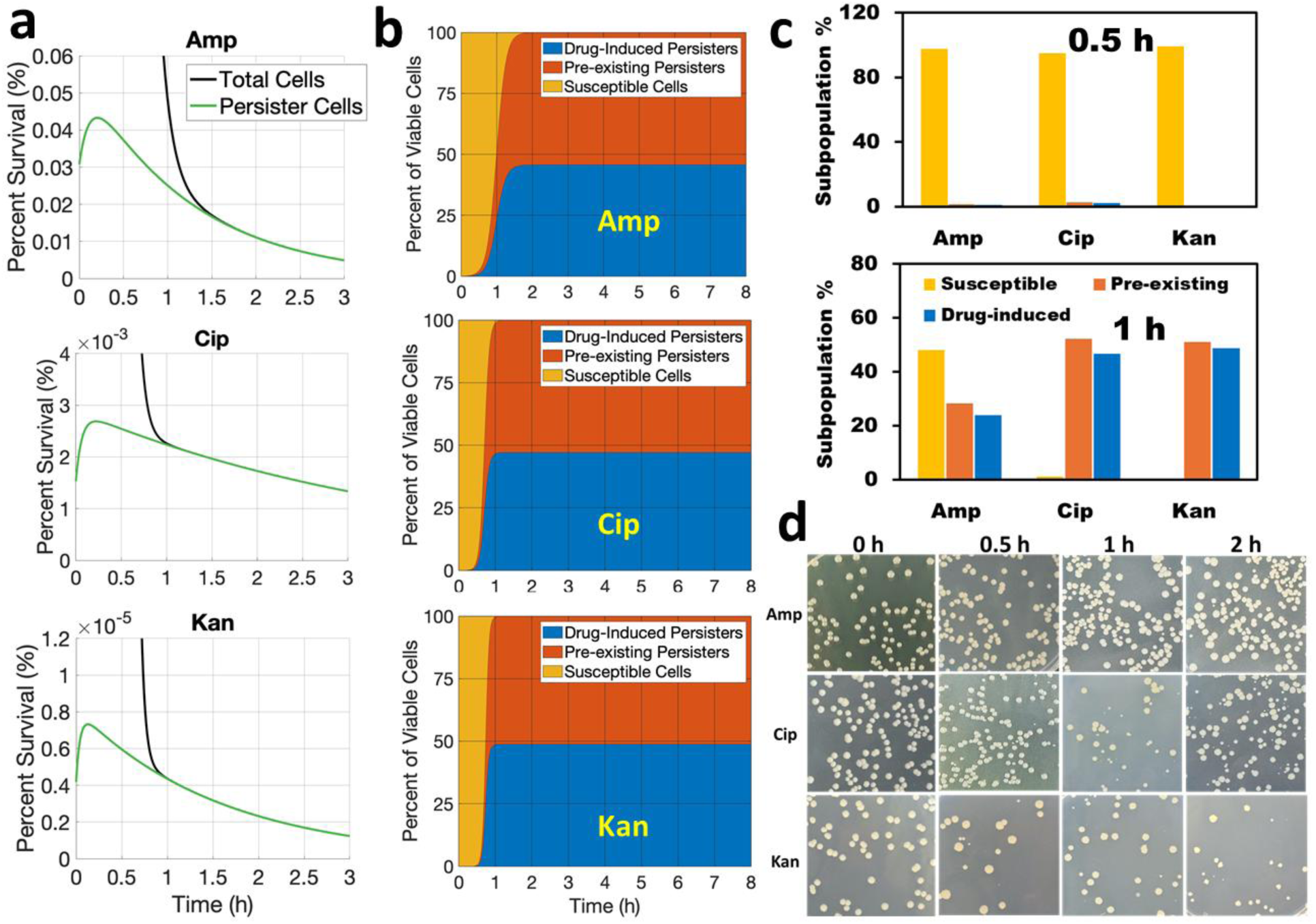
Dynamics of persister formation during antibiotic exposure. (a) Time-resolved dynamics of total population and persister subpopulations in 1,000-fold diluted cultures during antibiotic exposure, revealing rapid drug-induced persister formation. (b) Relative composition of susceptible cells, pre-existing persisters, and drug-induced persisters throughout the course of treatment. Note that the absolute numbers of surviving cells are markedly different across antibiotics. Simulations performed using the mean parameters presented in Fig. 3. (c) Relative fractions of susceptible cells, pre-existing persisters, and drug-induced persisters at 0.5 h and 1 h, extracted from model simulations shown in panel (b). (d) Appearance of small colonies after extended incubation, reflecting prolonged lag times and cellular damage following drug exposure.

Before treatment, distinct levels of pre-existing persisters were found across three antibiotics in the exponential-phase cultures (Fig. 5a). Upon drug exposure, the persister pool was quickly supplemented by newly induced persisters, producing an early peak of total persister abundance (Fig 5a). By ∼0.9-1.5 hours, nearly all surviving cells were persisters (Fig. 5a,b), of which ∼45–50% arose through drug-induced mechanisms, depending on the antibiotic (Fig. 5b) and seed dilution (Fig. 2d). These results indicate that bacteria do not passively succumb to antibiotic killing but actively respond to acute antibiotic stress by rapidly engaging defense and repair programs that enhance survival.

Population composition analysis further illustrates these dynamics (Fig. 5c). At 0.5 h, susceptible cells still dominated despite rapid persister formation. After 1 h of ampicillin treatment, the population comprised 48% susceptible cells, 28% pre-existing persisters, and 24% drug-induced persisters. In contrast, ciprofloxacin or kanamycin eliminated nearly all susceptible cells within 1 h, yielding survivors composed of 52% pre-existing and 47% induced persisters for ciprofloxacin, and 51% pre-existing and 49% induced persisters for kanamycin. Notably, the apparent balance between pre-existing and drug-induced persisters depends strongly on the seed dilution in SDTK assays; at high dilutions (e.g., 10,000-fold), survivors are dominated by induced persisters (Fig. S3b,c).

### Drug-specific differences reveal a spectrum of antibiotic persistence and hierarchical relationships among persister subsets

Across all three antibiotics, we consistently observed the emergence of small colonies in the viability plate count beginning with samples taken at ∼30 minutes of drug exposure, especially under ciprofloxacin and kanamycin treatment (Fig. 5d). These colonies typically became visible after an additional incubation for 2-3 days at room temperature following 20-h incubation at 37°C. Prolonged lag times and reduced growth are hallmarks of stressed, metabolically impaired, or partially damaged cells—phenotypes frequently observed during recovery from stationary phase, starvation, or other stressors ^16,36–38^. These observations illustrate that persister cells are highly heterogeneous. Upon antibiotic exposure, some cells encounter more severe damage than others and thus require longer recovery time before forming visible colonies on LB agar, resulting in variable colony sizes. This variability does not result from genetic mutations: when regrown, these persister cells formed normal-sized colonies indistinguishable from untreated wild type (Fig. S5).

Our findings that pre-existing kanamycin and ciprofloxacin persisters are orders of magnitude lower than ampicillin persisters (Fig. 3d, 5a) suggest that cells persistent to one antibiotic are not necessarily persistent to another. These distinct fractions of pre-existing persisters across drugs indicate substantial heterogeneity within the persister pool and drug-dependent thresholds for survival. Consistent with this interpretation, when multiple antibiotics were applied to the same exponential-phase culture, the resulting time-kill curves failed to extrapolate to a shared initial point (Fig. S4 a vs. Fig. S4 b and Fig. 6a), demonstrating that each drug reveals a different apparent “initial” persister level. These observations support a model in which persisters are distributed along a continuum of tolerance strengths, with different antibiotics inducing and revealing the population size of each drug’s persisters and their relative position along the tolerance spectrum.

**Fig. 6.**
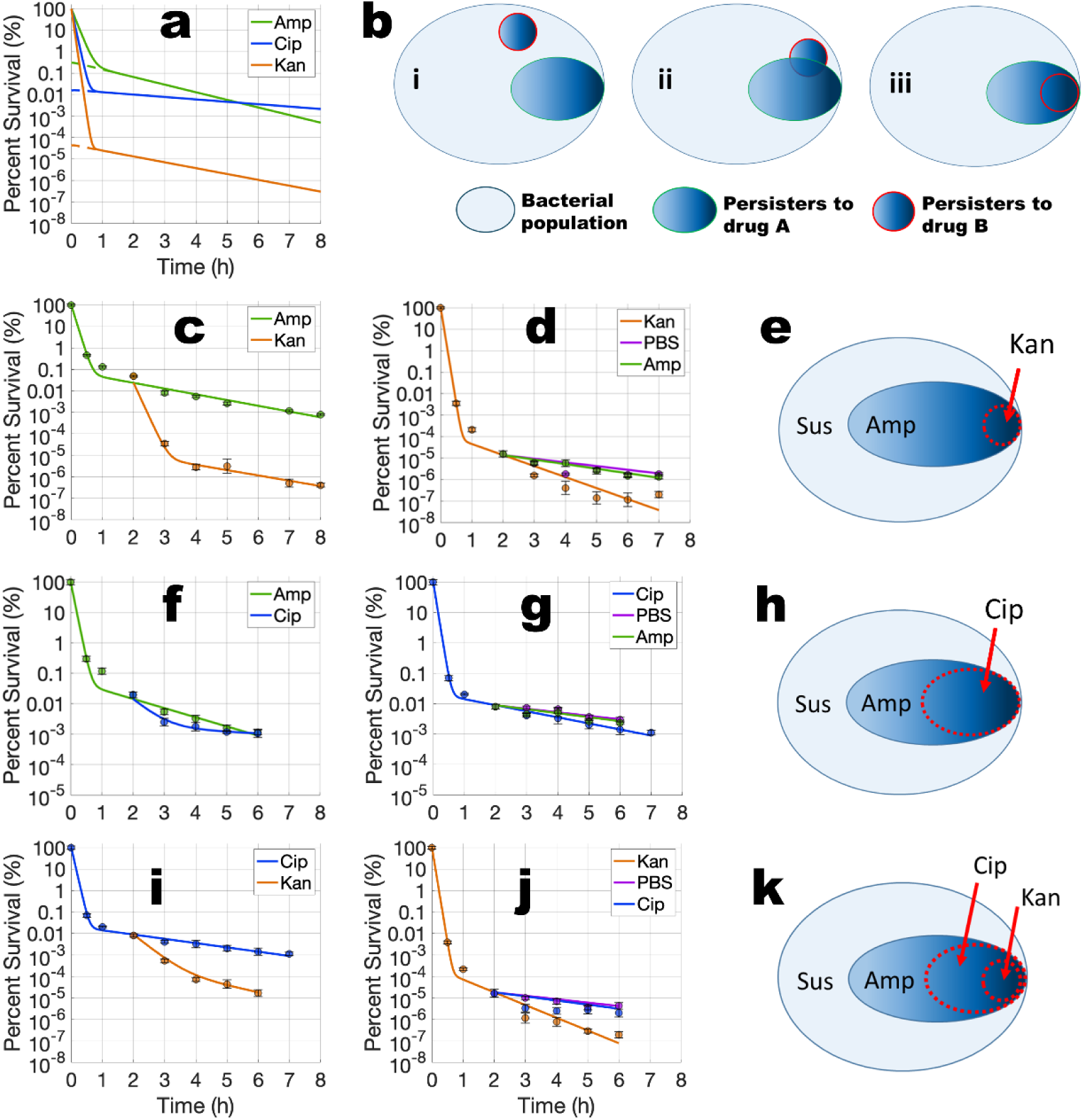
Hierarchical relationships and the spectrum of antibiotic persistence revealed by sequential antibiotic treatments. (a) Simulated dynamics of a bacterial culture exposed to ampicillin (Amp), ciprofloxacin (Cip) or kanamycin (Kan) using the averaged parameters measured from this work. Cultures from 100-fold seed dilution were chosen for the experiments and simulations because the effect of drug induction on the total persister pool is minimized under this dilution. (b) Graphic demonstration of the possible relationships between two persister subpopulations: (i) Two distinct subsets, (ii) Two partially overlapping subsets, (iii) One subset embedded within another subset. (c) Exponential-phase cultures were first treated with ampicillin (Amp) for 2 h, followed by exposure to either Kan or Amp after removal of the primary drug. (d) Cultures were first treated with Kan, followed by exposure to either Amp, Kan, or PBS after removal of the primary drug. PBS buffer was used as a negative control. (e) Proposed distribution and hierarchical relationship between the subsets of ampicillin persisters and kanamycin persisters. Susceptible cells (Sus), ampicillin persisters (Amp), and kanamycin persisters (Kan). (f) Cultures were first treated with Amp, followed by exposure to either Cip or Amp after removal of the primary drug. (g) Cultures were first treated with Cip, followed by exposure to either Amp, Cip, or PBS after removal of the primary drug. (h) Proposed distribution and hierarchical relationship between the subsets of ampicillin persisters and ciprofloxacin persisters. (i) Cultures were first treated with Cip, followed by exposure to either Cip or Kan after removal of the primary drug. (j) Cultures were first treated with Kan, followed by exposure to either Kan, Cip, or PBS after removal of the primary drug. (k) Proposed distribution and hierarchical relationship among the subsets of susceptible cells, ampicillin persisters, ciprofloxacin persisters, and kanamycin persisters. All cultures tested here were exponential-phase cultures initiated from a 100× dilution of the inoculum. Data represent means ± SD from four replicates across two independent experiments.

To resolve the relationship between the different persister subpopulations selected and induced by different antibiotics, we developed sequential drug treatment experiments. Exponential phase cultures were first exposed to one antibiotic for 2 hours, after which surviving persisters were washed, split, and challenged with either a second drug or the same drug again as a control. The experiments were then repeated with a reverse treatment order for the two antibiotics. This design enables us to determine whether persisters to one drug exhibit cross-persistence against another drug, thereby revealing the relative strength of drug-specific persistence.

Importantly, this approach also allows us to infer nested hierarchical relationships among persister subpopulations. Persisters selected and induced by two antibiotics can exhibit three possible relationships: (i) two non-overlapping subsets, (ii) partially overlapping subsets, or (iii) one subset fully embedded within the other (Fig. 6b), as proposed previously ^30^. Each relationship generates distinct predictions for sequential-drug treatments. When persister subsets are non-overlapping (relationship (i)), switching drugs results in rapid mutual elimination because each persister subset falls within the susceptible range of the other antibiotic. If the subsets partially overlap (relationship (ii)), each antibiotic should quickly eliminate the non-overlapping fraction of the other’s persisters. In contrast, when one drug’s persister subset is embedded within another (relationship (iii)), treatment with the former antibiotic should rapidly reduce the larger persister subset to a size comparable to the smaller, more persistent subset, whereas treatment with the latter should have little effect on the former’s persisters over a several-hour time window. Thus, sequential-drug treatments provide a direct means to resolve the hierarchical organization of persister subsets and their relative position along the persistence spectrum.

Sequential-drug treatments revealed a clear nested hierarchical relationship between the subsets of ampicillin and kanamycin persisters. Ampicillin persisters, which declined slowly under continued ampicillin exposure, were rapidly eliminated upon subsequent kanamycin treatment, producing biphasic killing dynamics nearly identical to kanamycin-only controls (Fig. 6c; Fig. 2a, Fig. 6a,d). This ∼3-log reduction demonstrates that most ampicillin persisters are not tolerant to kanamycin. In contrast, reversing the treatment order showed that ampicillin had minimal impact on kanamycin persisters, whose survival closely resembled PBS controls (Fig. 6d). The modest decline observed under ampicillin or PBS likely reflects post-stress death driven by delayed metabolic toxicity rather than direct ampicillin-mediated killing ^39^. Together, these results indicate that kanamycin persisters are cross-tolerant to ampicillin but not vice versa, establishing kanamycin persisters as a more persistent subpopulation.

Consistent with the embedded relationship between persister subsets, kanamycin persisters constitute a highly refractory population nested within the broader ampicillin-persister pool (Fig. 6e). Prolonged antibiotic exposure progressively enriched for increasingly tolerant cells, revealing an additional, slower-killing phase after extended treatment (24 h; Fig. S6). Persisters surviving long-term ampicillin exposure exhibited substantially reduced clearance upon kanamycin challenge compared with those surviving shorter ampicillin exposure (∼1-log versus ∼3-log reduction; Fig. S7a vs. Fig. 6c), indicating enrichment of a highly persistent subset corresponding to kanamycin persisters. The long-tail behavior has similarly been reported for lag times, recovery dynamics, and long-term killing kinetics ^40,41^, indicating heterogeneity along the persistence continuum. These and our observations support a nested structure in which extended antibiotic exposure selectively enriches for increasingly tolerant states along a persistence spectrum.

Sequential analyses further resolved the nested relationships among ampicillin, ciprofloxacin, and kanamycin persisters. Most ampicillin persisters were rapidly eliminated by ciprofloxacin, yielding biphasic killing dynamics, whereas prolonged ampicillin exposure enriched for a ciprofloxacin-tolerant subset (Fig. 6f; Fig. S7b). Conversely, ciprofloxacin persisters were largely unaffected by subsequent ampicillin treatment (Fig. 6g), indicating higher persistence. Finally, ciprofloxacin persisters were rapidly eliminated by kanamycin, while kanamycin persisters remained tolerant to ciprofloxacin (Fig. 6i,j).

Together, these results establish a nested persistence hierarchy in which kanamycin persisters represent the most tolerant subset, embedded within ciprofloxacin persisters, which are themselves a subset of ampicillin persisters (Fig. 6k). This framework supports a continuum model of persistence, explaining drug-specific cross-tolerance patterns and revealing how different antibiotics induce and select nested regions across the persistence spectrum.

## Discussion

In this study, we establish a quantitative framework by integrating kinetic modeling with serial-dilution time–kill (SDTK) assays. This approach enables accurate quantification of both pre-existing and drug-induced persisters within the same population—subpopulations that are otherwise operationally indistinguishable. Importantly, the SDTK framework resolves the challenge of measuring a biological variable (e.g., persister abundance) that is itself perturbed by the measurement process (e.g., time–kill assays). Using this strategy, we validate the drug-induced persister model and resolve population-level persister dynamics during antibiotic treatment without relying on specialized single-cell techniques. By circumventing key limitations of single-cell approaches, including low throughput, limited resolution, and dependence on reliable molecular markers, this framework enables quantitative analysis of extremely rare persister subpopulations.

Although the integrated SDTK framework allows simultaneous quantification of subpopulations, the accuracy of determining pre-existing persisters (P_0_) could be compromised when pre-existing persister fractions are extremely low. When cultures are extremely diluted, drug-induced persisters dominate (e.g., >99%), making the fractions of pre-existing persisters negligible and thus insensitive to model fitting. In practice, moderately diluted cultures provide the most reliable estimates. A simpler, coarse-grained alternative to SDTK quantification of persisters is to extrapolate the slow killing phase to time zero using minimally diluted cultures (e.g., ≤100-fold dilution, in which drug-induced persister formation varied by <15%) (Fig. 2d). This simplified approach provides better estimates of persister cells than the single time-point CFU counts but should be considered an upper-bound estimate of pre-existing persister subpopulation size.

Our drug-induced kinetic model focuses on the first 7-8 hours of antibiotic exposure, during which biphasic killing predominates, but additional killing phases can emerge with prolonged treatment ^10^. Indeed, extending exposure to 24 hours reveals a multi-phase pattern consisting of rapid killing of susceptible cells, slower decay of most persisters, and an even slower phase corresponding to ultra-persistent cells (Fig. S6). These highly refractory cells represent fewer than 10^-8^ of the initial population and are several orders of magnitude rarer than the dominating persisters captured by our model. Therefore, excluding these ultra-rare subpopulations does not affect our model’s accuracy. The biphasic framework provides a robust and reliable description of drug-induced persister dynamics over our experimental timescale.

### Why do different antibiotics confer varying killing and drug-induced persister formation rates?

Differences in persister frequency and killing dynamics across antibiotics can be largely explained by their distinct modes of action and metabolic dependencies. Ampicillin, a β-lactam whose activity strongly depends on metabolism, is ineffective against slow-growing or dormant cells ^42^. Growth arrest under acute ampicillin stress is sufficient to convert cells into persisters, producing the largest pre-existing persister fraction and the rapidest induction of new persisters (Figs. 2,3). Ciprofloxacin, in contrast, remains effective against non-growing cells through lethal DNA damage ^42^, resulting in smaller pre-existing persister fractions. Survival requires activation of additional stress-response pathways, such as SOS-mediated DNA repair, yielding a smaller but more refractory persister subset. Kanamycin, an aminoglycoside, irreversibly disrupts ribosomes ^43^, cell membranes and the cytoplasm ^44,45^. With the least metabolic dependence among the three antibiotics tested ^42^, kanamycin rapidly eliminates susceptible cells and most Amp and Cip persisters, leaving the rarest but most tolerant survivors. These differences together with differential cellular responses to antibiotics ^46^ explain why ampicillin, ciprofloxacin, and kanamycin select and generate distinct persistence phenotypes.

Interestingly, despite being less potent than ciprofloxacin against susceptible cells, ampicillin kills persisters faster (greater *k₂*) than ciprofloxacin (Fig. 3b,c). This apparent paradox likely reflects differences in the persister states generated: ampicillin readily selects and produces new persisters, but these cells are under a less protected physiological state and are therefore more easily killed, whereas ciprofloxacin generates fewer persisters, but they are considerably more recalcitrant. Differences in target accessibility and binding reversibility may further contribute: ampicillin irreversibly inhibits penicillin-binding proteins on cell surfaces ^47^, whereas ciprofloxacin must enter the cell before binding to DNA-associated enzymes, resulting in relatively weak and reversible drug-enzyme-DNA complexes when drug is removed ^48,49^.

### A Drug-Induced Persistence Spectrum (DIPS) model to explain persister dynamics and heterogeneity

Our results demonstrate that persistence is heterogeneous and spans across a spectrum of tolerance. Sequential-drug treatments and extended time-kill analyses reveal nested hierarchies of persister subpopulations (Fig. 6k). Prolonged exposure progressively enriches for increasingly persistent cells with slower killing kinetics (Fig. S6 and S7). This graded heterogeneity is supported by single-cell observations that persisters vary widely in growth rate, lag time, metabolic activity, and stress responses ^22,23,40,50–52^. The slow-growing colonies of different sizes observed after prolonged incubation in our assays and others ^16,36^ likely reflects these graded states and recovery capacities. Even persisters with the same persistence can arise from different combinations of target inactivation, cellular damages, and stress-triggered defense and repair programs.

Here, we propose a Drug-Induced Persistence Spectrum (DIPS) model describing the formation and dynamics of heterogeneous persisters during antibiotic treatment (Fig. 7). Antibiotic persisters are not created equal. At the onset, the stationary-phase inoculum exhibits a wide range of dormancy depths and lag times ^40,52^. Upon transfer to fresh medium, most cells rapidly resume growth, whereas a small heterogeneous fraction remains dormant, metabolically slow, or in an adaptive state, constituting the pre-existing persister pool. Once exposed to antibiotics, exponentially growing cells undergo acute stress and extensive injury, and the majority are rapidly eliminated with a high killing rate constant (*k₁*) before protective pathways can be activated. A minority, however, successfully activate sufficient stress-induced responses to repair DNA, protein and/or membrane damage, surviving the primary attack and drug-induced reactive species ^39,53–55^. These cells thus transition into persisters at a drug-specific formation rate (*f₁*), acquiring various persistence levels. When drug-inflicted damages exceed the capacity of defense and repair systems, both pre-existing and drug-induced persisters are ultimately eliminated with a slower killing rate constant (*k₂*).

**Figure 7.**
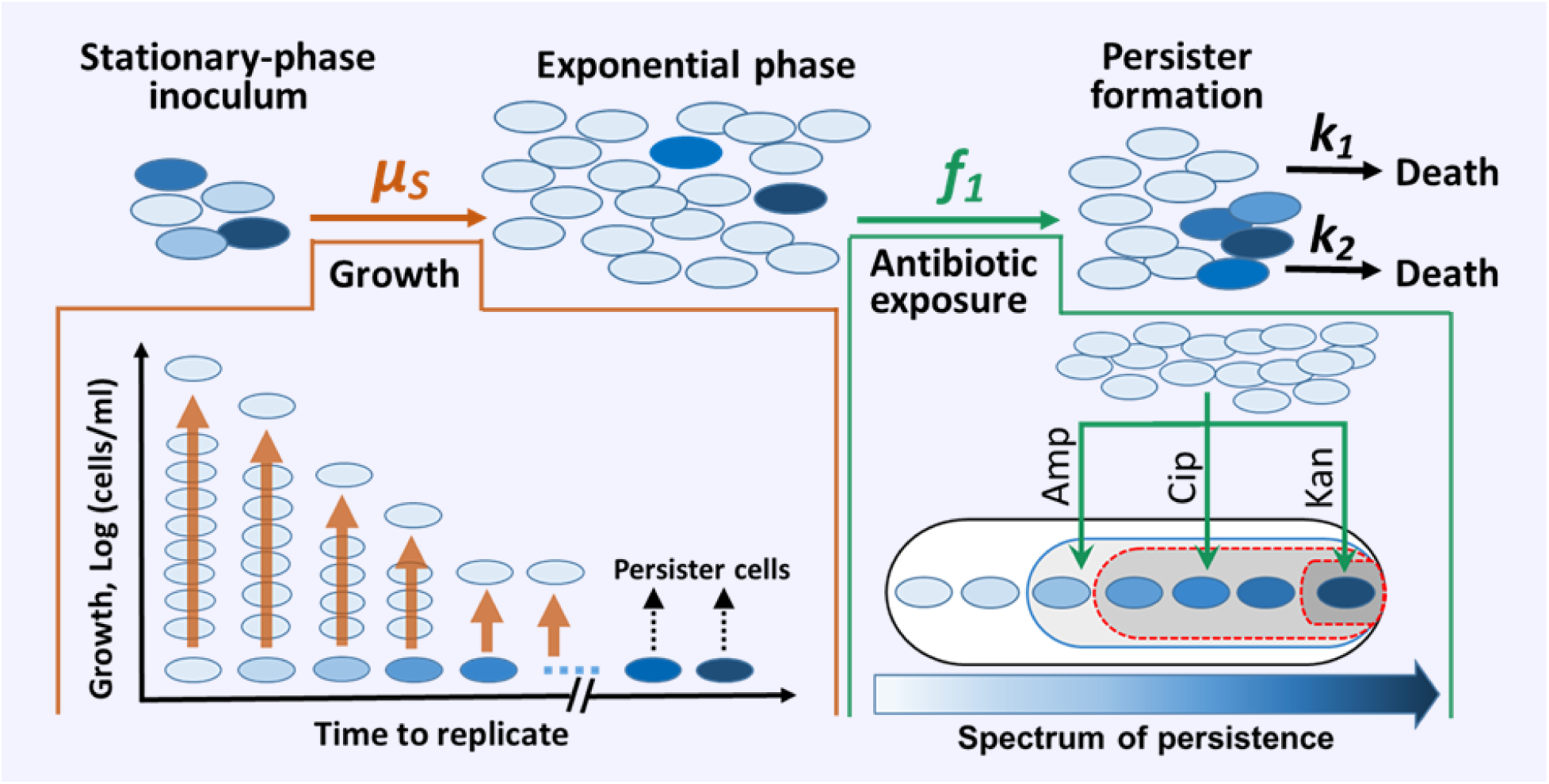
A proposed model for the dynamics of antibiotic persistence. Upon inoculation into fresh medium, most cells rapidly resume growth (light color) and enter the exponential phase, while a small subpopulation, the pre-existing persisters (darker color), remains dormant and tolerant to antibiotics. The revival time of cells follows a power-law distribution with most cells replicating quickly (light color) and a few remaining dormant as pre-existing persisters (dark color). The darkness of color denotes the strength of persistence, with darker color indicating higher persistence. During antibiotic exposure, actively growing cells die rapidly (rate constant *k₁*), whereas a minority activate stress-induced protective responses (e.g., ROS defenses, and/or SOS repair responses) and transition into newly formed persisters (rate constant *f₁*). Antibiotic persisters are not created equal: different drugs induce different degrees of persistence, and even the same drug can trigger heterogeneity of persistence. Kanamycin selects and generates a small subset of highly persistent cells, while ampicillin selects and induces a relatively weak but large persister pool, with ciprofloxacin’s effects being intermediate. Both pre-existing and drug-induced persisters exhibit a spectrum of persistence levels and can be ultimately killed (killing rate constant *k₂*) when antibiotic-induced damage exceeds their repair capacity.

Ampicillin selects and generates the largest but weakest persister subset, ciprofloxacin induces an intermediate subset, and kanamycin yields a rare, highly refractory subset. These subsets form a nested hierarchy across the persistence spectrum, with kanamycin persisters embedded within ciprofloxacin persisters, which are themselves nested within the broader ampicillin-persister pool (Fig 7). The spectrum of persistence yields sequential long-term killing dynamics: rapid elimination of susceptible cells followed by progressively slower decay of increasingly tolerant persisters. The resulting long-tailed kinetics can be described by power-law or Weibull model ^40,41,56^, while short-term dynamics are well approximated by biphasic exponential decay.

This DIPS framework reconciles divergent observations in persister profiles across antibiotics ^12,30,57^. More broadly, it can be generalized into a “stress-induced-persistence-spectrum” model, as environmental stresses such as heat, low pH, oxidative stress, hazardous chemicals, and starvation also trigger antibiotic persistence ^7,10,58^. These stresses cause varying degrees of cellular damage and induce different levels of defense and repair mechanisms in bacterial cells, resulting in heterogeneity in persistence. Because antibiotics differ in their propensity to induce and enrich tolerant subpopulations, treatment can select highly persistent cells even without genetic resistance. Incorporating drug-induced persistence profiles alongside MIC evaluations may therefore guide antibiotic choice and improve strategies to limit persister emergence and accelerate clearance.

## Conclusion

In summary, this work establishes a simple yet powerful framework that integrates SDTK assays with kinetic modeling to accurately resolve the population dynamics of susceptible cells, pre-existing persisters, and drug-induced persisters. Applying this approach, we show that antibiotics differ not only in killing efficacy but also in their capacity to induce persisters, generating persister subpopulations that vary markedly in size and tolerance. Our framework further enables the identification of genetic determinants that specifically govern pre-existing persistence and/or persister formation during treatment. Through systematic sequential-drug treatments, we demonstrate hierarchical relationships among antibiotic-specific persister subsets and propose a DIPS model, demonstrating that persistence is heterogeneous and distributed along a tolerance spectrum from which each antibiotic induces and selects distinct subsets. Kanamycin persisters, although exceedingly rare, represent the most refractory extreme, whereas ciprofloxacin persisters occupy an intermediate position within the broader ampicillin persister population. This work provides a broadly applicable strategy for dissecting drug tolerance in other systems, including fungal pathogens and cancer cells associated with treatment failure and relapse.

## Methods and Materials

### Model development

In time-kill assays, antibiotic persistence can be characterized by biphasic killing kinetics, with a rapid elimination phase of susceptible cells followed by a slow killing phase of persisters. In these assays, stationary-phase cultures are diluted into pre-warmed fresh medium and grown to exponential phase before antibiotic addition. At the time of drug exposure, the population is assumed to consist of two major subpopulations: susceptible cells (*S*) and persister cells (*P*). Spontaneously formed persisters arising during exponential growth are presumed negligible compared with the pre-existing persisters inherited from stationary phase. Thus, although more heavily diluted cultures require longer times to reach the same density, the number of spontaneous persisters in all dilutions is assumed to remain negligible (discussed in the main text).

We further assume that persisters carried over from the stationary-phase inoculum do not replicate during cultivation, despite potentially retaining low metabolic activity ^28^. Although single-cell studies have reported that a small fraction of persisters can slowly grow prior to antibiotic exposure ^22,23,59^, this behavior does not affect the validity and accuracy of our model or the interpretation of the SDTK experiments. Any potential replication of pre-existing persisters would occur proportionally across serially diluted cultures and therefore preserve dilution scaling predicted by a pre-existing-only model (Fig. 1b). Moreover, only a minority of persister cells are capable of growth, and their replication rates are substantially slow ^16,23,59^. The effect of persister replication would therefore be minimal compared with the number of persisters generated from the orders-of-magnitude larger susceptible population during antibiotic exposure. Thus, the drug-induced persister model remains robust, even without explicitly incorporating into our framework the small fraction of growing persisters or spontaneous persister formation.

Both *P* and *S* subpopulations are killed independently, with rate constants *k₁* for susceptible cells and *k₂* for persisters. The population dynamics during antibiotic exposure are therefore described as:

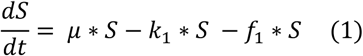

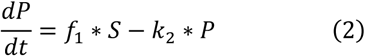

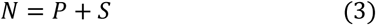

Here, *f₁*represents the rate constant of drug-induced persister formation. The classical pre-existing-persister model is a special case of this framework, *f₁ = 0*, in which no additional persisters form during treatment. All parameters and variables, including *f₁*, *k₁*, *k₂*, *S₀* (initial susceptible population), and *P₀*(initial persister population), were obtained from experimental measurements and model fitting. We assume that bacterial growth halts immediately upon drug supplement (*μ = 0*).

### Strains and growth conditions

The wild-type E. coli strain MG1655 was used for most experiments. Two knockout strains, including *Δppk* and *ΔrelAΔspoT*, each derived from MG1655, were kindly provided by Gray lab ^32,60^ and used to assess the roles of polyphosphate kinase (PPK) and the ppGpp pathway in persister formation. Cultures were grown in Luria–Bertani (LB; Miller formulation) medium with or without antibiotics. Three bactericidal antibiotics with distinct modes of action were examined: ampicillin (Amp) targeting cell wall synthesis; ciprofloxacin (Cip) targeting DNA gyrase; and kanamycin (Kan) targeting the ribosome. Drug stocks were freshly prepared and stored at −20 °C. Amp solutions were stored at −80 °C and used within two months. Unless otherwise noted, antibiotics were applied at 10× MIC. For turbitdostat experiments, a continuous-culture device (Chi.Bio) ^61^ was used to maintain exponential-phase growth (OD₆₀₀ ≈ 0.6) for ∼12 h prior to initiating SDTK assays.

### Serial-dilution time-kill (SDTK) assays

Single colonies were grown overnight in LB, diluted 1:100,000 into fresh LB medium, and cultured for 18 h to generate seed cultures. We used this two-step cultivation to minimize experimental variations and the potential age effect on antibiotic persistence ^62^. These were then serially diluted 100×, 1,000×, or 10,000× into pre-warmed fresh LB medium. Cultures were grown to early exponential phase with OD₆₀₀ ≈ 0.6 (∼2–3 × 10⁸ cells ml⁻¹) before antibiotic addition. Samples were collected at defined intervals for 6–8 h. After sampling, antibiotics were removed by washing, and cells were plated on antibiotic-free LB agar plates. Plates were incubated at 37 °C for 20 h and then held at room temperature for an additional 2 days to allow small colonies to emerge. For kanamycin-treated samples where killing was substantially faster, cultures were concentrated prior to plating, and plates were kept at room temperature for another three days for small colonies to appear before CFU enumeration.

### MIC determination

Minimum inhibitory concentrations (MICs) were measured for the wild type, *ΔrelAΔspoT*, and *ΔppK* strains following previously described protocols ^63^ with slight modification. Each of the three antibiotics (ampicillin, ciprofloxacin, and kanamycin), was serially diluted 2-fold in LB medium in 96-well microtiter plates. Overnight cultures were then diluted into fresh LB medium and grown to exponential phase (∼ 2-3 h), then further diluted in pre-warmed LB medium and inoculated into the wells containing the antibiotic dilutions at the same initial cell density (OD_600_= ∼0.02). After incubation at 37 °C overnight, OD₆₀₀ values were measured using a plate reader (Molecular Devices, Inc.) to assess bacterial growth. The MIC for each drug-strain combination was defined as the lowest antibiotic concentration at which no increase in OD₆₀₀ was observed.

### Sequential antibiotic treatment

Seed cultures prepared as described above were diluted 100-fold into fresh medium and grown to an OD₆₀₀ of 0.6 before the addition of a primary antibiotic. Unless otherwise mentioned, two hours after exposure, the drug was removed by centrifuging the cultures at 37 °C and resuspending the pellets in pre-warmed LB medium supplemented either with a second antibiotic (for sequential treatment) or with the same antibiotic (for continuous single-drug treatment). The cultures were then centrifuged and resuspended once more in the same pre-warmed LB medium to ensure complete removal of the primary drug. As a control, after removal of the primary antibiotic, cultures were washed and resuspended in pre-warmed phosphate-buffered saline (PBS). Samples were collected at defined time intervals, and surviving cells were enumerated by CFU plating. The extended-exposure experiments were performed as described above, except that the initial antibiotic exposure was prolonged for the durations specified in the Results. Unless otherwise mentioned, we used the cultures grown from the 100-fold diluted seed culture to ensure that most persisters tested were from the pre-existing fractions and to minimize the confounding effects of drug-induced persisters.

### Model fitting

Kinetic models were simulated and fitted to experimental SDTK data using MATLAB (version 2024b, MathWorks Inc.). Model dynamics were simulated using the “ode45()” solver. For fitting, parameters k_1_ and k_2_ were fitted to the first and second linear regions, respectively, of the log-transformed viable cell counts using the first-order (linear) polyfit() function. Then, lsqnonlin() performed simultaneous fitting of P_0_ for the least diluted culture (e.g., 100×) and *f_1_* for all cultures. *P_0_* values for the more dilute cultures were set by the corresponding dilution factors. For turbidostat-Amp fits, the initial persister percent, *P_0_* %, was assumed to be 8×10^-5^ based on simulations (Fig S2d, Table S1) of a previously published spontaneous persister formation model ^16^. For sequential treatment data fitting (Fig. 6), *f_1_* was assumed to be zero because the contribution of drug-induced persisters is minimal in those experiments due to their low seed dilution (100-fold).

### Statistical methods

All statistical analyses were performed using GraphPad Prism (Version 10.2.3). Group differences were assessed by one-way ANOVA, followed by Tukey’s Honest Significant Difference (HSD) post hoc test, with significance set at α < 0.05. When assumptions of normality were violated, the nonparametric Kruskal–Wallis test was applied. When normality was met but variances were unequal, Brown–Forsythe or Welch ANOVA was used, followed by Dunnett’s post hoc test for multiple comparisons. For two-group comparisons with unequal variances, Welch’s t-test was employed.

## Data availability

All data used in this study are available in the main manuscript and the Supplementary material files; raw data are available upon request.

## Code availability

All code files and related data used in this study are available in the GitHub repository: https://github.com/schultz-lab/persisters.

## Acknowledgements

We thank Michal Gray at the University of Alabama at Birmingham for kindly providing the *E coli MG1655 Δppk* and *ΔrelAΔspoT* strains. We thank Wolfgang L. Weber, Iam A. Cucho, and Eunice A. Antwi for their assistance in collecting experimental data.

## Contributions

**Y.D. and R.S.** conceived the project. **Y.D.** designed the experiments. **Y.D. and D.R.B.** developed the models. **Y.D., H.E.M., and K.E.** performed the experiments and collected the data. **D.R.B.** performed model simulations and data fitting. **D.R.B. and Y.D.** analyzed the data and prepared the figures. **R.S. and D.S.** secured the funding and resources. **Y.D.** wrote the manuscript. **Y.D., D.R.B., D.S., and R.S.** revised the manuscript. All authors read and approved the final manuscript.

## Supplementary Information

**Supplementary Figure S1.**
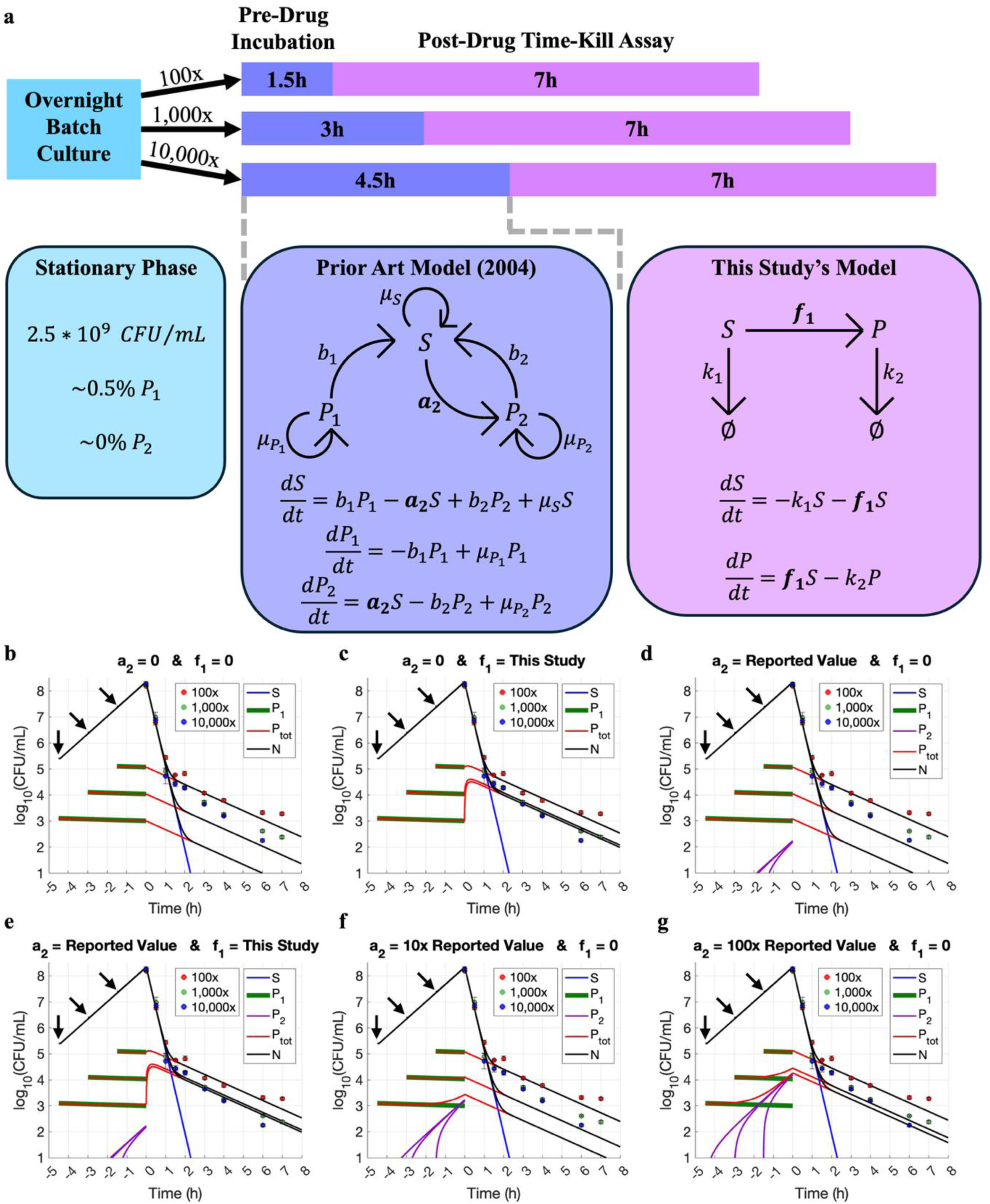
Spontaneous persister formation does not explain the observed time-kill data from overnight batch cultures. (a) A schematic representation of the experimental setup and the associated kinetic modeling for each step. In the prior art model incorporating spontaneous persister formation (Balaban *et al.* 2004), *P_1_* represents pre-existing persisters from the stationary phase inoculum and *P_2_* represents spontaneous persisters that arise from susceptible cells at a rate *a_2_*. In this figure, we used our experimental data and model fitting to compare the pre-existing persister model (b), our drug-induced persister model (c), the prior art spontaneous-persister-formation model (d) and the combined model of the latter two (e), modeling the persister dynamics from the beginning of inoculation until the end of the time-kill assay in each case. Results show that our model faithfully captures experimental data (c) while the prior art model does not (d). Incorporation of spontaneous persister formation is not necessary because the contribution of spontaneous persisters is negligible compared to the pre-existing and drug-induced persisters (e). To explore the possibility for the prior art model to fit our experimental data, we arbitrarily increased *a_2_* for the wild-type strain to biologically implausible values by 10-fold (f) or 100-fold (g). We found that the spontaneous persister formation rate must increase by at least 100-fold to achieve a reasonable fit (g). In other words, only when the spontaneous persister fraction becomes orders of magnitude larger, dominating the overall persister subpopulation, can the spontaneous-persister-formation model begin to fit the observed SDTK data. Under normal conditions, however, spontaneous persisters in the wild-type strain typically constitute only a small fraction of the total persister population because the spontaneous persister formation rate for the wild-type strain is expected to be extremely low, as reported by Balaban *et al* (2004). In panels (b-g), the curves for the 100-, 1,000-, and 10,000-fold diluted cultures are time-shifted to align the start of drug treatment at t=0. The inoculation times of these diluted pre-drug cultures are marked by arrows, with the 10,000-fold diluted culture starting at 4.5 hours before drug exposure (-4.5 h). The experimental data are from Fig. 2a, an ampicillin SDTK assay performed on batch-seeded wild-type *E coli* cultures.

**Supplementary Figure S2.**
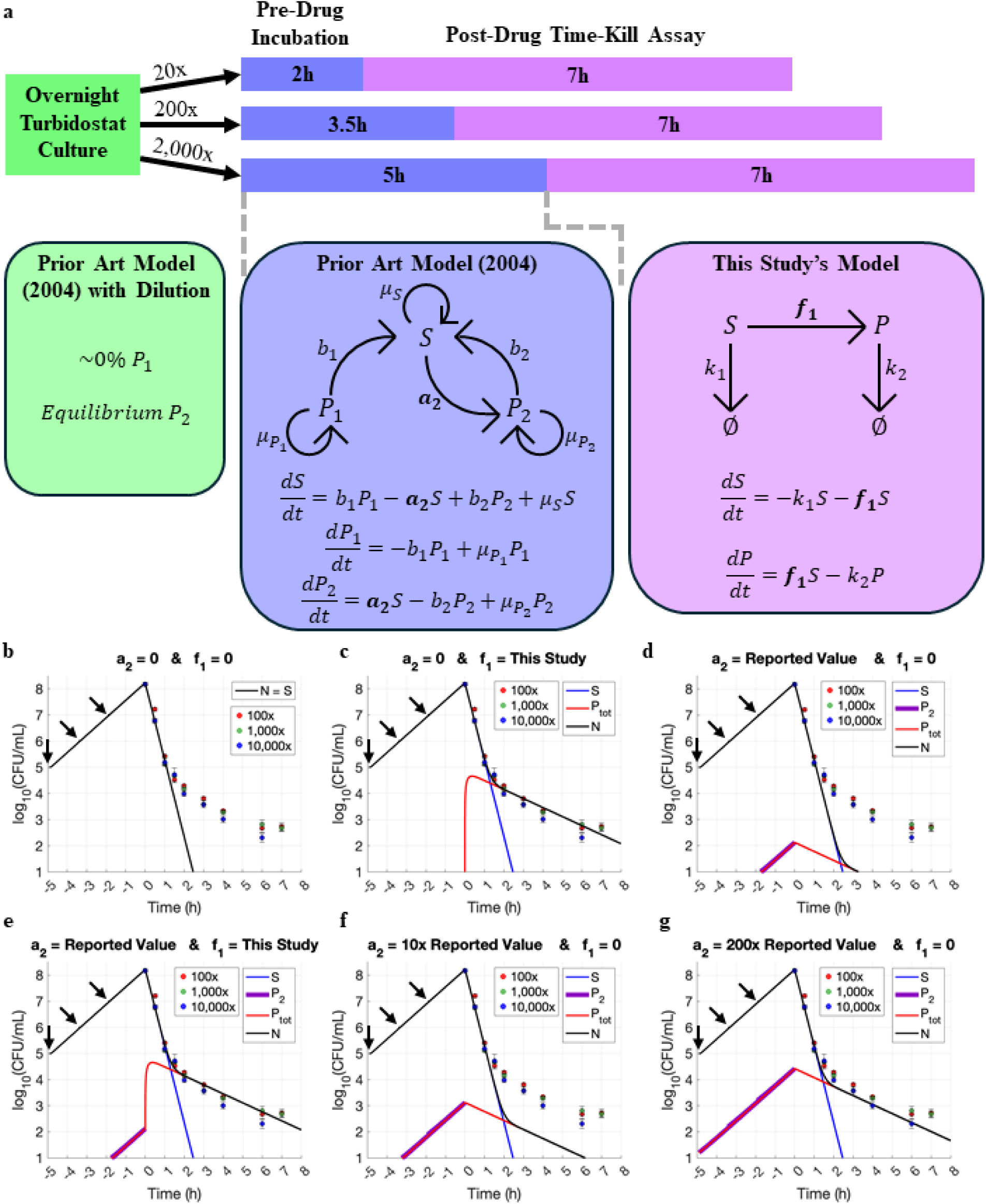
Spontaneous persister formation does not explain the observed time-kill data from turbidostat experiments. This figure largely parallels Supplementary Figure S1 but with an overnight turbidostat culture instead of an overnight batch culture. (a) A schematic representation of the experimental procedure and the associated kinetic modeling for each step. The prior art model is the spontaneous-persister-formation model developed by Balaban *et. al.* (2004). The pre-existing persister model (b) cannot fit our experimental data because the continuous turbidostatic cultivation removes virtually all pre-existing persisters from the seed culture. Simulation results show that our drug-induced persister model faithfully captures all experimental data (c) while the prior spontaneous-persister-formation model does not (d). Incorporation of spontaneous persister formation is not necessary because the contribution of spontaneous persisters is negligible compared to the drug-induced persisters (e). To explore the possibility for the prior art model to fit our experimental data, we arbitrarily increased the spontaneous persister formation rate (*a_2_*) for the wild-type strain to biologically implausible values by 10-fold (f) or 200-fold (g). We found that the spontaneous persister formation rate must increase by at least 200-fold to achieve a reasonable fit (g). In other words, only when the spontaneous persister formation rate becomes orders of magnitude larger than reported value can the spontaneous-persister-formation model begin to capture the observed experimental data. In panels b-g, the curves for the 20-, 200-, and 2000-fold diluted cultures are time-shifted to align the start of drug treatment at t = 0. The inoculation times of these diluted pre-drug cultures are marked by arrows, with the 2000-fold diluted culture starting at 5 hours before drug exposure (-5 h). The experimental data for fitting are from Fig. 2c, an ampicillin SDTK assay performed on turbidostat-seeded wild-type *E coli* cultures.

**Supplementary Fig. S3.**
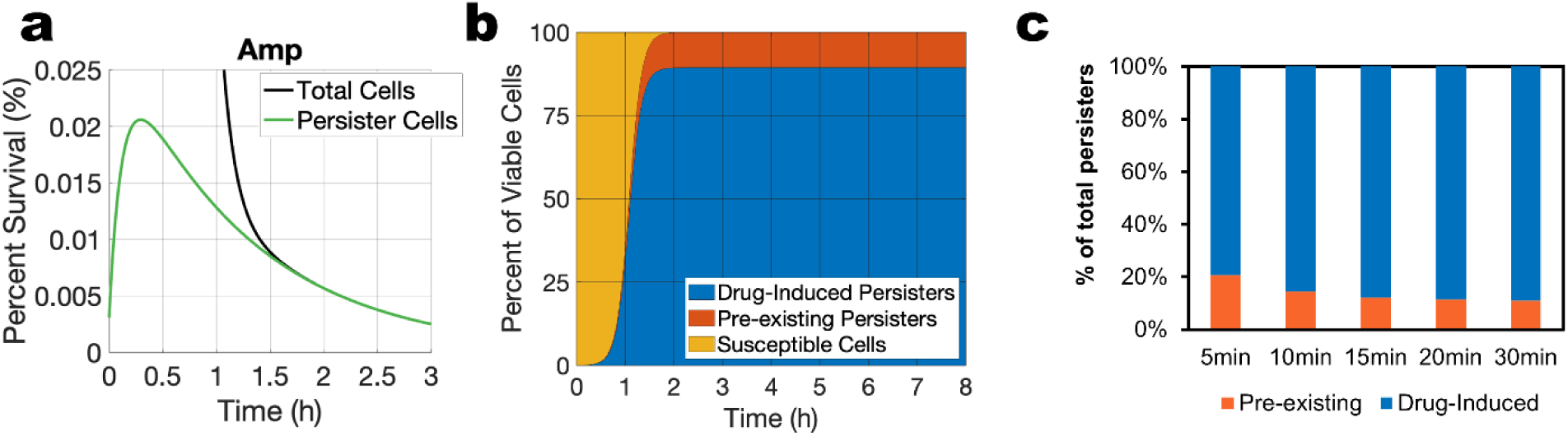
Model simulation of persister dynamics in cultures derived from 10,000-fold seed dilution. (a) Dynamics of persisters and total cells exposed to ampicillin. (b) Relative composition of surviving cells over 8 hours of treatment with ampicillin. (c) Relative fractions of pre-existing and drug-induced persisters in the first 30 min upon ampicillin exposure. The relative ratio reaches a steady state within 20 min of exposure.

**Supplementary Fig. S4.**
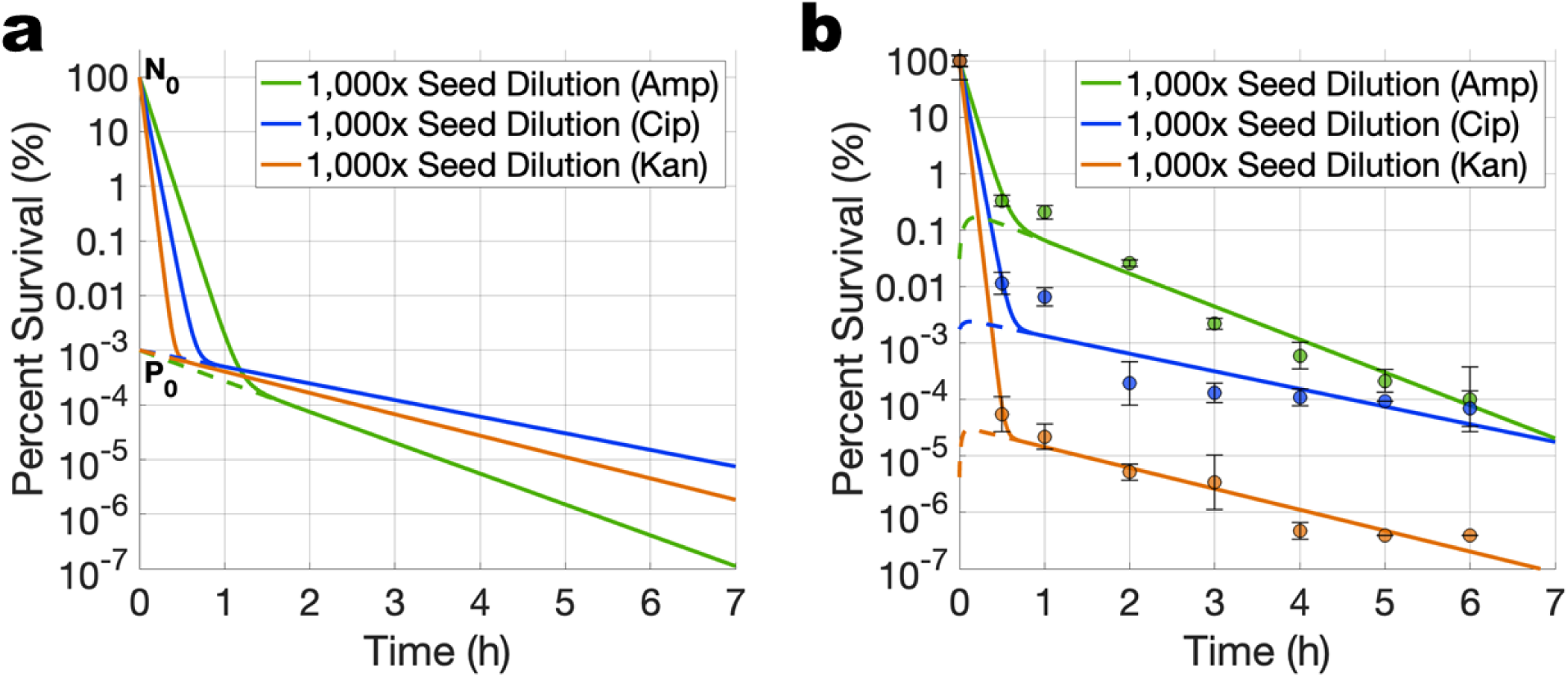
Same exponential-phase cultures were treated by different antibiotics. (a) A static persister model assumes that the same pre-existing subpopulation is persistent to different antibiotics. (b) Experimental data showing the same cultures treated by ampicillin (Amp), ciprofloxacin (Cip), and kanamycin (Kan), each at 10x MICs. The data support the drug-induced persister model and rule out the static model in which the same persister subpopulation is assumed to tolerate different antibiotics. Data points represent means ± SD from four replicates across two independent experiments.

**Supplementary Fig. S5.**
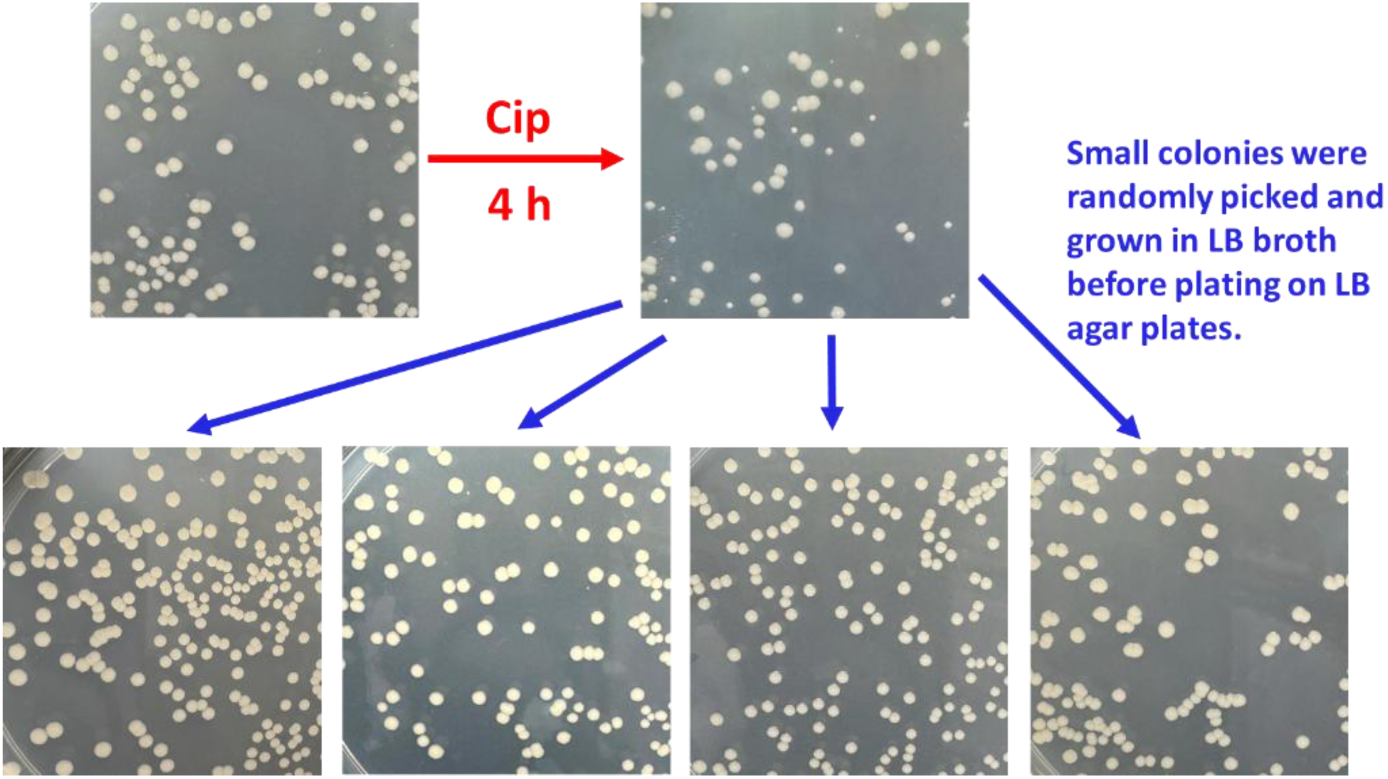
The small colonies observed after antibiotic treatment do not result from genetic mutations. When isolated and regrown, the small colonies emerging from ciprofloxacin treatment display colony size and morphology indistinguishable from the wild type prior to antibiotic exposure. Exponential-phase cultures were treated with ciprofloxacin (Cip) for 4 hours, after which persister cells were washed and plated on LB agar. Plates were incubated at 37 °C for 20 hours and then at room temperature for an additional 1-2 days to allow small colonies to emerge. These small colonies were randomly picked, grown in LB broth to exponential phase, and replated on LB agar to assess their colony sizes and morphology. In all cases, the regrown colonies matched wild-type characteristics, confirming that the initial small-colony phenotype was transient and not mutation-driven.

**Fig. S6.**
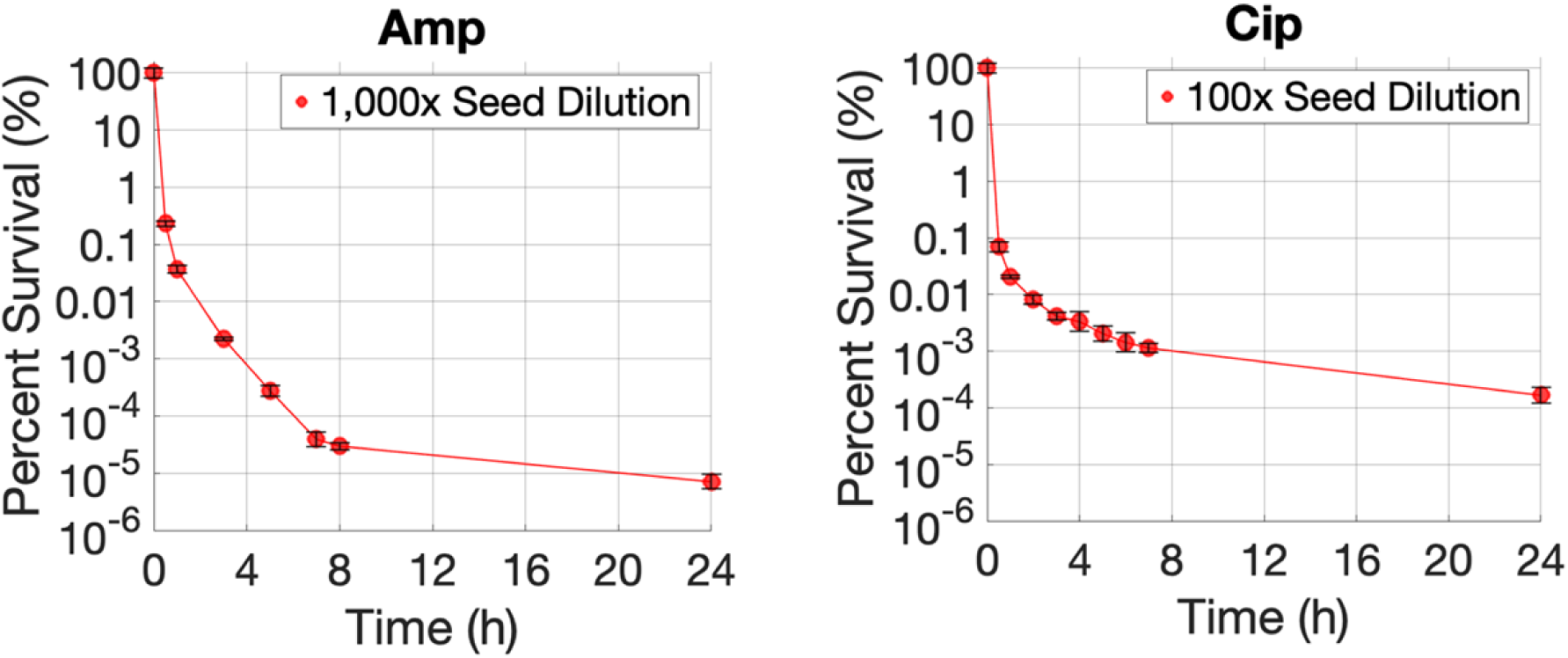
Extended time-kill assays over a 24-hour period. (a) Time-kill kinetics of cultures treated with ampicillin, initiated from a 1,000-fold diluted inoculum. (b) Time-kill kinetics of cultures treated with ciprofloxacin, initiated from a 100-fold diluted inoculum. The killing dynamics exhibit at least three distinct phases. During the first 7∼8 hours, two major phases are evident: rapid killing of susceptible cells, followed by slower killing of the persister subpopulation. From around 8 to 24 hours, a third phase becomes apparent, corresponding to the slowest elimination of an extremely small subpopulation of highly persistent cells. These “high-persistence” subsets were orders of magnitude smaller than the major persister fraction (∼0.1%). Because this subset is negligible relative to the primary killing phases, our drug-induced kinetic model does not incorporate this ultra-rare persistence component. Data represent means ± SD from four replicates across two independent experiments.

**Fig. S7.**
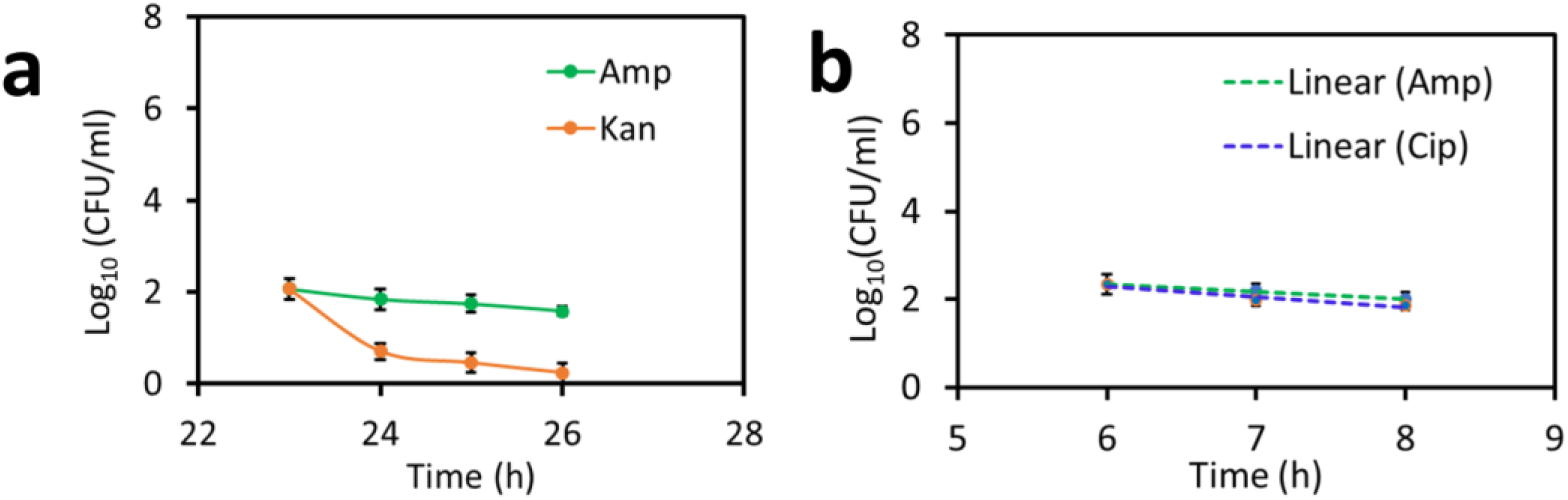
Sequential drug treatments for cultures with extended exposure. (a) Bacteria were treated with ampicillin (Amp) for 23 h before exposed to kanamycin (Kan). (b) Bacteria were treated with Amp for 6 h before exposed to ciprofloxacin (Cip) or Amp. Data represent are means ± SD from four replicates across two independent experiments.

**Table S1:** Relative contributions of spontaneous and drug-induced persister formation during SDTK assays in turbidostatic experiments. The sequence of models used for Table S1 is same as shown in Fig S2a: the prior art spontaneous-persister-formation model in turbidostatic culture (pre-existing stationary-phase persisters get diluted out and spontaneous persisters form) and in pre-drug batch culture (spontaneous persister formation), and our drug-induced model during drug exposure (drug-induced persister formation). We assume that the turbidostatic culture removes all stationary-phase pre-existing persisters by continuous dilution. Therefore, all of the persisters observed in the time-kill assay are either spontaneously formed or drug-induced. In this special case, we can independently fit the spontaneous persister formation rate, *a_2_*, and the drug-induced persister formation rate, *f_1_*. As a result, given the value of any one parameter among *a₂*, *f₁*, and *P₀* (the fraction of persisters immediately before drug exposure), the other two parameters/variables can be uniquely determined. Specifically, *a₂*and *P₀* are constrained by the prior spontaneous-persister-formation model of Balaban *et. al.* (2004), such that specifying one fixes the other. The value of *f_1_* is set by *P_0_*, or vice versa, by fitting SDTK data with our kinetic model. The table shows hypothetical scenarios; a 650-fold increase in the reported *a_2_* value would be needed to explain the experimental data without *f_1_*. In this case, the persister fraction reached an unrealistical value of 0.052% in the exponential phase, which is not plausible for the wild-type strain under this continuously diluted conditions. In contrast, only a 1.004 multiplier on our fitted value of *f_1_* would be needed to explain the experimental data without *a_2_*. Using the previously measured value for *a_2_*, over 99.8% of all persisters are drug-induced after 1 hour of drug exposure. The experimental dataset fitted to generate this table is the same as shown in Fig. 2C and Supplementary Fig S2, a turbidostat-seeded ampicillin SDTK assay.

| $P_0$<br>(% of tot. cells) | $a_2$ (1/h) | $a_2/a_2^0$<br>* | Spontaneous<br>$P$ (% of $P_{tot}$) | $f_1$ (1/h) | $f_1/f_1^0$<br>** | Drug-Induced<br>$P$ (% of $P_{tot}$) |
| --- | --- | --- | --- | --- | --- | --- |
| 0.052*** | $7.8 \times 10^{-4}$ | 650 | 100 | 0 | 0 | 0 |
| 0.01 | $1.5 \times 10^{-4}$ | 125 | 24.2 | $1.87 \times 10^{-3}$ | 0.76 | 75.8 |
| $10^{-3}$ | $1.5 \times 10^{-5}$ | 12.5 | 2.42 | $2.41 \times 10^{-3}$ | 0.98 | 97.58 |
| **$8 \times 10^{-5}$ (Fig. 2c)** | **$1.2 \times 10^{-6}$** | **1** | **0.19** | **$2.46 \times 10^{-3}$** | **1** | **99.81** |
| $10^{-5}$ | $1.5 \times 10^{-7}$ | 0.125 | 0.02 | $2.47 \times 10^{-3}$ | 1.004 | 99.98 |
| 0 | 0 | 0 | 0 | $2.47 \times 10^{-3}$ | 1.004 | 100 |

\* $a_2^0$ is the spontaneous persister formation rate constant value ( $a_2 = 1.2 \times 10^{-6} \text{ h}^{-1}$ ) previously measured for the wild-type *E. coli* strain exposed to ampicillin (Balaban *et al.* 2004).
\*\* $f_1^0$ is our fitted value of persister formation rate constant ( $f_1 = 2.46 \times 10^{-3} \text{ h}^{-1}$ ) for bacteria exposed to ampicillin.
\*\*\* This $P_0$ value (the persister fraction before drug exposure) was determined by extrapolating the experimental data of the second killing phase, assuming $f_1 = 0$ . Other $P_0$ values in column 1 are hypothetical scenarios except for the bolded row. The bolded text denotes that the simulation results were obtained using experimentally measured values of $f_1$ and $a_2$ .

## References

1. Murray, C. J. et al. Global burden of bacterial antimicrobial resistance in 2019: a systematic analysis. Lancet 399, 629–655 (2022).

2. Fisher, R. A., Gollan, B. & Helaine, S. Persistent bacterial infections and persister cells. Nat. Rev. Microbiol. 15, 453–464 (2017).

3. Bakkeren, E., Diard, M. & Hardt, W.-D. Evolutionary causes and consequences of bacterial antibiotic persistence. Nat. Rev. Microbiol. 18, 479–490 (2020).

4. Lewis, K. Persister Cells. Annu. Rev. Microbiol. 64, 357–372 (2010).

5. Windels, E. M. et al. Bacterial persistence promotes the evolution of antibiotic resistance by increasing survival and mutation rates. ISME J. 13, 1239–1251 (2019).

6. Bigger, J. Treatment of staphylococcal infections with penicillin by intermittent sterilization. Lancet 244, 497–500 (1944).

7. Keren, I., Kaldalu, N., Spoering, A., Wang, Y. & Lewis, K. Persister cells and tolerance to antimicrobials. FEMS Microbiol. Lett. 230, 13–18 (2004).

8. Rotem, E. et al. Regulation of phenotypic variability by a threshold-based mechanism underlies bacterial persistence. Proc. Natl. Acad. Sci. 107, 12541–12546 (2010).

9. Levin, B. R., Concepción-Acevedo, J. & Udekwu, K. I. Persistence: a copacetic and parsimonious hypothesis for the existence of non-inherited resistance to antibiotics. Curr. Opin. Microbiol. 21, 18–21 (2014).

10. Balaban, N. Q. et al. Definitions and guidelines for research on antibiotic persistence. Nat. Rev. Microbiol. 17, 441–448 (2019).

11. Song, S. & Wood, T. K. ppGpp ribosome dimerization model for bacterial persister formation and resuscitation. Biochem. Biophys. Res. Commun. 523, 281–286 (2020).

12. Harms, A., Maisonneuve, E. & Gerdes, K. Mechanisms of bacterial persistence during stress and antibiotic exposure. Science 354, (2016).

13. Zhang, Y. Persisters, persistent infections and the Yin–Yang model. Emerg. Microbes Infect. 3, 1–10 (2014).

14. Podlesek, Z. & Žgur Bertok, D. The DNA Damage Inducible SOS Response Is a Key Player in the Generation of Bacterial Persister Cells and Population Wide Tolerance. Front. Microbiol. 11, (2020).

15. Niu, H., Gu, J. & Zhang, Y. Bacterial persisters: molecular mechanisms and therapeutic development. Signal Transduct. Target. Ther. 9, 174 (2024).

16. Balaban, N. Q., Merrin, J., Chait, R., Kowalik, L. & Leibler, S. Bacterial persistence as a phenotypic switch. Science 305, 1622–1625 (2004).

17. Manuse, S. et al. Bacterial persisters are a stochastically formed subpopulation of low-energy cells. PLoS Biol. 19, (2021).

18. Johnson, P. J. T. & Levin, B. R. Pharmacodynamics, Population Dynamics, and the Evolution of Persistence in Staphylococcus aureus. PLoS Genet. 9, e1003123 (2013).

19. Dörr, T., Lewis, K. & Vulić, M. SOS Response Induces Persistence to Fluoroquinolones in Escherichia coli. PLoS Genet. 5, e1000760 (2009).

20. Dörr, T., Vulić, M. & Lewis, K. Ciprofloxacin Causes Persister Formation by Inducing the TisB toxin in Escherichia coli. PLoS Biol. 8, e1000317 (2010).

21. Greve, N. B., Slotved, H.-C., Olsen, J. E. & Thomsen, L. E. Identification of antibiotic induced persister cells in Streptococcus agalactiae. PLoS One 19, e0303271 (2024).

22. Orman, M. A. & Brynildsen, M. P. Dormancy Is Not Necessary or Sufficient for Bacterial Persistence. Antimicrob. Agents Chemother. 57, 3230–3239 (2013).

23. Goormaghtigh, F. & Van Melderen, L. Single-cell imaging and characterization of Escherichia coli persister cells to ofloxacin in exponential cultures. Sci. Adv. 5, (2019).

24. El Meouche, I., Jain, P., Jolly, M. K. & Capp, J.-P. Drug tolerance and persistence in bacteria, fungi and cancer cells: Role of non-genetic heterogeneity. Transl. Oncol. 49, 102069 (2024).

25. Russo, M. et al. Cancer drug-tolerant persister cells: from biological questions to clinical opportunities. Nat. Rev. Cancer 24, 694–717 (2024).

26. Sharma, S. V. et al. A Chromatin-Mediated Reversible Drug-Tolerant State in Cancer Cell Subpopulations. Cell 141, 69–80 (2010).

27. Rosenberg, A. et al. Antifungal tolerance is a subpopulation effect distinct from resistance and is associated with persistent candidemia. Nat. Commun. 9, 2470 (2018).

28. Rahman, K. M. T. et al. Rethinking dormancy: Antibiotic persisters are metabolically active, non-growing cells. Int. J. Antimicrob. Agents 65, 107386 (2025).

29. Manina, G. & McKinney, J. D. A Single-Cell Perspective on Non-Growing but Metabolically Active (NGMA) Bacteria. in Pathogenesis of Mycobacterium tuberculosis and its Interaction with the Host Organism 135–161 (2013). doi:10.1007/82_2013_333.

30. Shultis, M. W., Mulholland, C. V. & Berney, M. Are all antibiotic persisters created equal? Front. Cell. Infect. Microbiol. 12, (2022).

31. Gray, M. J. & Jakob, U. Oxidative stress protection by polyphosphate—new roles for an old player. Curr. Opin. Microbiol. 24, 1–6 (2015).

32. Hamm, C. W. & Gray, M. J. Inorganic polyphosphate and the stringent response coordinately control cell division and cell morphology in Escherichia coli. MBio 16, (2025).

33. Fritsch, V. N. et al. The alarmone (p)ppGpp confers tolerance to oxidative stress during the stationary phase by maintenance of redox and iron homeostasis in Staphylococcus aureus. Free Radic. Biol. Med. 161, 351–364 (2020).

34. Qi, W., Jonker, M. J., de Leeuw, W., Brul, S. & ter Kuile, B. H. Role of RelA-synthesized (p)ppGpp and ROS-induced mutagenesis in de novo acquisition of antibiotic resistance in E. coli. iScience 27, 109579 (2024).

35. Bergum, O. E. T. et al. SOS genes are rapidly induced while translesion synthesis polymerase activity is temporally regulated. Front. Microbiol. 15, (2024).

36. Vulin, C., Leimer, N., Huemer, M., Ackermann, M. & Zinkernagel, A. S. Prolonged bacterial lag time results in small colony variants that represent a sub-population of persisters. Nat. Commun. 9, (2018).

37. Guillier, L., Pardon, P. & Augustin, J.-C. Influence of Stress on Individual Lag Time Distributions of Listeria monocytogenes. Appl. Environ. Microbiol. 71, 2940–2948 (2005).

38. Levin-Reisman, I. et al. Automated imaging with ScanLag reveals previously undetectable bacterial growth phenotypes. Nat. Methods 7, 737–739 (2010).

39. Hong, Y., Zeng, J., Wang, X., Drlica, K. & Zhao, X. Post-stress bacterial cell death mediated by reactive oxygen species. Proc. Natl. Acad. Sci. 116, 10064–10071 (2019).

40. Şimşek, E. & Kim, M. Power-law tail in lag time distribution underlies bacterial persistence. Proc. Natl. Acad. Sci. 116, 17635–17640 (2019).

41. Rebelo, J. S., Domingues, C. P. F., Monteiro, F., Nogueira, T. & Dionisio, F. Bacterial persistence is essential for susceptible cell survival in indirect resistance, mainly for lower cell densities. PLoS One 16, e0246500 (2021).

42. Zheng, E. J., Stokes, J. M. & Collins, J. J. Eradicating Bacterial Persisters with Combinations of Strongly and Weakly Metabolism-Dependent Antibiotics. Cell Chem. Biol. 27, 1544–1552.e3 (2020).

43. Davis, B. D. Mechanism of bactericidal action of aminoglycosides. Microbiological Reviews vol. 51 341–350 (1987).

44. Wohlgemuth, I. et al. Translation error clusters induced by aminoglycoside antibiotics. Nat. Commun. 12, 1830 (2021).

45. Wong, F. et al. Cytoplasmic condensation induced by membrane damage is associated with antibiotic lethality. Nat. Commun. 12, (2021).

46. Mattiello, S. P., Barth, V. C., Scaria, J., Ferreira, C. A. S. & Oliveira, S. D. Fluoroquinolone and beta-lactam antimicrobials induce different transcriptome profiles in Salmonella enterica persister cells. Sci. Rep. 13, 18696 (2023).

47. Beadle, B. M., Nicholas, R. A. & Shoichet, B. K. Interaction energies between β-lactam antibiotics and E. coli penicillin-binding protein 5 by reversible thermal denaturation. Protein Sci. 10, 1254–1259 (2001).

48. Maglica, Ž., Özdemir, E. & McKinney, J. D. Single-cell tracking reveals antibiotic-induced changes in mycobacterial energy metabolism. MBio 6, e02236–14 (2015).

49. Mustaev, A. et al. Fluoroquinolone-gyrase-DNA complexes: two modes of drug binding. J. Biol. Chem. 289, 12300–12 (2014).

50. Ma, P. et al. Bacterial droplet-based single-cell RNA-seq reveals antibiotic-associated heterogeneous cellular states. Cell 186, 877–891.e14 (2023).

51. Stevanovic, M., Teuber Carvalho, J. P., Bittihn, P. & Schultz, D. Dynamical model of antibiotic responses linking expression of resistance genes to metabolism explains emergence of heterogeneity during drug exposures. Phys. Biol. 21, 036002 (2024).

52. Pu, Y. et al. ATP-Dependent Dynamic Protein Aggregation Regulates Bacterial Dormancy Depth Critical for Antibiotic Tolerance. Mol. Cell 73, 143–156.e4 (2019).

53. Wong, F. et al. Reactive metabolic byproducts contribute to antibiotic lethality under anaerobic conditions. Mol. Cell 82, 3499–3512.e10 (2022).

54. Kohanski, M. A., Dwyer, D. J., Hayete, B., Lawrence, C. A. & Collins, J. J. A Common Mechanism of Cellular Death Induced by Bactericidal Antibiotics. Cell 130, 797–810 (2007).

55. Schultz, D., Palmer, A. C. & Kishony, R. Regulatory Dynamics Determine Cell Fate following Abrupt Antibiotic Exposure. Cell Syst. 5, 509–517.e3 (2017).

56. Peleg, M. Modeling the dynamic kinetics of microbial disinfection with dissipating chemical agents—a theoretical investigation. Appl. Microbiol. Biotechnol. 105, 539–549 (2021).

57. Brauner, A., Fridman, O., Gefen, O. & Balaban, N. Q. Distinguishing between resistance, tolerance and persistence to antibiotic treatment. Nat. Rev. Microbiol. 14, 320–30 (2016).

58. Cotten, K. L. & Davis, K. M. Bacterial heterogeneity and antibiotic persistence: bacterial mechanisms utilized in the host environment. Microbiol. Mol. Biol. Rev. 87, (2023).

59. Umetani, M. et al. Observation of persister cell histories reveals diverse modes of survival in antibiotic persistence. Elife 14, (2025).

60. Gray, M. J. Inorganic Polyphosphate Accumulation in Escherichia coli Is Regulated by DksA but Not by (p) ppGpp. (2019).

61. Steel, H., Habgood, R., Kelly, C. L. & Papachristodoulou, A. In situ characterisation and manipulation of biological systems with Chi.Bio. PLOS Biol. 18, e3000794 (2020).

62. Luidalepp, H., Jõers, A., Kaldalu, N. & Tenson, T. Age of inoculum strongly influences persister frequency and can mask effects of mutations implicated in altered persistence. J. Bacteriol. 193, 3598–3605 (2011).

63. Barnes V, L., Heithoff, D. M., Mahan, S. P., House, J. K. & Mahan, M. J. Antimicrobial susceptibility testing to evaluate minimum inhibitory concentration values of clinically relevant antibiotics. STAR Protoc. 4, 102512 (2023).

